# Intragenic conflict in phylogenomic datasets

**DOI:** 10.1101/2020.04.13.038133

**Authors:** Stephen A. Smith, Nathanael Walker-Hale, Joseph F. Walker

**Author notes:** Equal contribution.

## Abstract

Most phylogenetic analyses assume that a single evolutionary history underlies one gene. However, both biological processes and errors in dataset assembly can violate this assumption causing intragenic conflict. The extent to which this conflict is present in empirical datasets is not well documented. However, if common, it would have far-reaching implications for phylogenetic analyses. Here, we examined several large phylogenomic datasets from diverse taxa using a fast and simple method to identify well supported intragenic conflict. We found conflict to be highly variable between datasets, from 1% to more than 92% of genes investigated. To better characterize patterns of conflict, we analyzed four genes with no obvious data assembly errors in more detail. Analyses on simulated data highlighted that alignment error may be one major source of conflict. Whether as part of data analysis pipelines or in order to explore potential biologically compelling intragenic processes, analyses of within gene signal should become common. The method presented here provides a relatively fast means for identifying conflicts that is agnostic to the generating process. Datasets identified with high intragenic conflict may either have significant errors in dataset assembly or represent conflict generated by biological processes. Conflicts that are the result of error should be identified and discarded or corrected. For those conflicts that are the result of biological processes, these analyses contribute to the growing consensus that, similar to genomes, genes themselves may exhibit multiple conflicting evolutionary histories across the tree of life.

## Introduction

The sequencing and analysis of whole genomes, transcriptomes, and thousands of individual genes has illustrated that, throughout the tree of life, genomes are a composite of evolutionary histories. This heterogeneity is a source of biological insight (Mendes et al. 2019) as well as a source of computational and analytical complexity (Kosakovsky Pond et al. 2006a,b; Boussau et al. 2013; Smith et al. 2015). Though heterogeneity is found across the tree of life, researchers have primarily focused on conflict among trees inferred using individual genes, in many cases limited to combined exons, and the inferred species tree.

Driven primarily by an interest in identifying recombination break points, over the past two decades researchers have examined some sources of heterogeneous topological signal within single-gene alignments. To facilitate this, several methods have been developed (e.g., Husmeier and McGuire 2003, Hobolth et al. 2007, Suchard et al. 2002, Allman et al. 2017, Ane 2011, Bossau et al. 2009, Kosakovsky Pond et al 2006a, 2006b). While recombination undoubtedly plays a large role in population genetics, at the species level, the impact of recombination has on phylogenetic inference remains debated (Edwards 2009, Lanier and Knowles, 2012; Wu 2013). Scornavacca and Galtier (2017) showed that exons within the same gene may present different genealogies, a pattern confirmed by Mendes et al. (2019). Despite this proliferation of methods, many are not tractable for phylogenomic datasets and/or assume the conflict arose from a biological process.

While biological processes can introduce phylogenetic heterogeneity within gene sequence alignments, given the dataset size and automated nature of genomic analyses, systematic error may also contribute to intragenic phylogenetic conflict in empirical datasets. These alignment and/or assembly errors have become more common as datasets have increased in both taxon sampling and regions of the genome analyzed. The volume of data has made automation a requirement of any phylogenomic pipeline. Inevitably, errors make their way into alignments and, if not filtered, can lead to phylogenetic conflict and downstream errors (Song et al. 2012; Gatesy and Springer, 2013; Brown and Thomson 2017, Walker et al. 2018).

Regardless of the source, incorrectly modeling intragenic conflict will result in inaccurate phylogenies, biased branch lengths, erroneous selection analyses, and can influence downstream analyses. Importantly, how common mixed signal is within genes is still unknown. Whether due to computationally taxing methods or because researchers assume that errors will be overwhelmed by the signal from hundreds of other genes (but see Brown and Thomson 2017; Shen et al. 2017; Walker et al 2018), analyses of intragenic conflict are not common. Here, instead of modeling the biological processes that generate conflicting signal (e.g., recombination), we assess violations of the underlying assumptions of phylogenetic reconstruction (e.g., mixed signal). This allows for faster analyses on phylogenomic datasets. We examine several empirical datasets using a sliding window approach and characterize the extent to which single gene-regions show evidence for conflicting histories across a broad range of phylogenetic datasets.

## Materials and Methods

### Empirical datasets

We gathered a broad sampling of nucleotide datasets across the tree of life. Many of these include a mix of exons, introns, and other genetic elements. For simplicity, when we refer to gene, we mean a locus or set of genetic elements but not necessarily a complete or protein-coding gene. This conforms to typical naming conventions such as “gene tree”, which may or may not refer to the tree of a coding sequence or complete gene. The datasets examined here included those designed to analyzed contentious relationships broadly across mammals: MAM1 consisting of 10,259 genes (Chen et al. 2017); MAM2 consisting of 424 genes(Song et al. 2012 as refined by Mirarab et al. 2014); and MAM3 consisting of 183 UCE loci (McCormack et al. 2012). We analyzed an insect dataset (BUGS) focused on analyzing relationships in the Strepsiptera consisting of 4,485 genes (Niehuis et al. 2012). We examined three vertebrate datasets: VERT, consisting of 1,113 genes (Wang et al. 2013); FISH, a dataset assembled to understand the relationships among ray finned fish, consisting of 1,105 genes (Hughes et al. 2018); and FROG, an dataset of frogs, consisting of 95 genes (Feng et al. 2017). We analyzed three plant datasets: PLAN, a dataset generated to investigate land plant evolution, consisting of 852 genes (Wickett et al. 2014); MOS(Mitochondria [M],Nuclear [N],Plastome[P]), a dataset generated to analyze moss ordinal relationships, consisting of three datasets of 40, 105, and 82 genes (Liu et al. 2019); and CARN, a carnivorous plant dataset, that consisted of 1,237 genes (Walker et al. 2017). Finally, we gathered a fungal dataset (FUNG) consisting of 2,256 genes (Pizzaro et al. 2018). As a result of missing data in segments of alignments, length of alignment not permitting the division into multiple segments, along with other factors that would prevent species relationships from being inferred from segments of the genes, the number of genes analyzed was often lower than the number from the original study (Table 2).

**Table 1.** Results from simulation analyses of differing alignment lengths simulated under different models with different within-gene conflicts. The percentage of replicates where the ML tree for the whole alignment was incorrectly inferred are indicated by “bad ML”; an asterisk indicates that not all these replicates overlapped with cases where multiple trees were detected.

| *Type* | *Description* | *% genes w. conflicts* |
| --- | --- | --- |
| No conflict | 25 t 3000 bp JC | 0% |
|  | 50 t 3000 bp JC | 0% |
|  | 25 t 1500 bp JC | 0% |
|  | 25 t 3000 bp diff. GTR | 0% |
|  | 25 t 3000 bp random GTR | 1% |
| Rate shift | 25 t 3000 bp random GTR diff. t height | 0% |
| Indel | 25 t 3000 bp random GTR indel equal | 1% |
|  | 25 t 3000 bp random GTR indel equal – MAFFT | 10% (10% bad ML*) |
|  | 25 t 3000 bp random GTR indel equal – FSA | 0% (2% bad ML) |
|  | 25 t 3000 bp random GTR indel equal – PRANK | 6% (10% bad ML*) |
|  | 25 t 3000 bp random GTR indel diff. 1 | 1% |
|  | 25 t 3000 bp random GTR indel diff. 1 – MAFFT | 11% (18% bad ML*) |
|  | 25 t 3000 bp random GTR indel diff. 1 – FSA | 1% (6% bad ML*) |
|  | 25 t 3000 bp random GTR indel diff. 1 – PRANK | 18% (19% bad ML*) |
|  | 25 t 3000 bp random GTR indel diff. 2 | 2% |
|  | 25 t 3000 bp random GTR indel diff. 2 – MAFFT | 12% (14% bad ML*) |
|  | 25 t 3000 bp random GTR indel diff. 2 – FSA | 2% (2% bad ML) |
|  | 25 t 3000 bp random GTR indel diff. 2 – PRANK | 18% (11% bad ML*) |
| Conflict | 25 t 3000 bp random GTR 200 bp conf. | 19.5% |
|  | 25 t 3000 bp random GTR 500 bp conf. | 68.4% |
|  | 25 t 3000 bp random GTR 1000 bp conf. | 98% |
|  | 25 t 1500 bp random GTR 200 bp conf. | 51% |

**Table 2.** Results from analyses of individual datasets including the number of taxa and number of genes in the original study, the number of genes long enough to analyze, and the proportion of those analyzed with conflict.

| ***Dataset*** | ***# of taxa*** | ***# of regions*** | ***# of analyzed regions*** | ***% w. conflict*** |
| --- | --- | --- | --- | --- |
| MAM1 (Chen et al. 2017) | 22 | 10,259 | 3,666 | 6.5% |
| MAM2 (Song et al. 2012) | 37 | 424 | 293 | 27.7% |
| MAM3 (McCormack et al. 2012) | 29 | 183 | 1 | 100% |
| VERT (Wang et al. 2013) | 12 | 1,113 | 551 | 2.7% |
| BUGS (Niehuis et al. 2012) | 13 | 4,485 | 467 | 1.9% |
| FISH (Hughes et al. 2018) | 303 | 1105 | 118 | 92.4% |
| FROG (Feng et al. 2017) | 164 | 95 | 40 | 80% |
| FUNG (Pizzaro et al. 2018) | 51 | 2,256 | 1,750 | 37.2% |
| CARN (Walker et al. 2017) | 13 | 1,237 | 343 | 0.6% |
| MOSM (Liu et al. 2019) | 134 | 40 | 11 | 100% |
| MOSN (Liu et al. 2019) | 134 | 105 | 81 | 85.2% |
| MOSP (Liu et al. 2019) | 134 | 82 | 23 | 87% |
| PLAN (Wickett et al. 2014) | 103 | 852 | 160 | 16.3% |

### Identification of multiple trees within a gene

For each dataset, we examined the conflicting signal inferred from contiguous segments of each gene. For most datasets we used 1000 bp for the segment length to increase phylogenetic signal per segment and reduce gap only taxa within segments. However, for datasets with generally shorter alignments including FISH, FROGS, and MOSS, we considered a segment length of 500 bp. Phylogenetic trees were calculated for the entire gene and for each gene segment using IQ-TREE (Nguyen, 2015), the GTR+Γ model of molecular evolution, 1000 Ultra-FastBootstrap (UFB) replicates (Hoang et al. 2018), and SH-aLRT (Guindon et al 2010) analyses. While the UFB has been shown to be conservative with at a cutoff of 95% (Hoang et al. 2018), with any relationship whose support value is below that is inferred to have low support, our simulations demonstrated that this still generated support (BS => 95%) for some poorly supported relationships. To be more conservative, we therefore calculated both SH-aLRT (with a cutoff of 80% used) and UFB. We then compared the segment trees to the maximum likelihood tree for the entire geneand gene segments found to contain conflicting signal, after considering the support cutoff, for any given relationship were extracted to be examined using the sliding window approach described below (Fig. 1A). These analyses were conducted with the python program phynd (https://github.com/FePhyFoFum/phynd).

**Figure 1.**
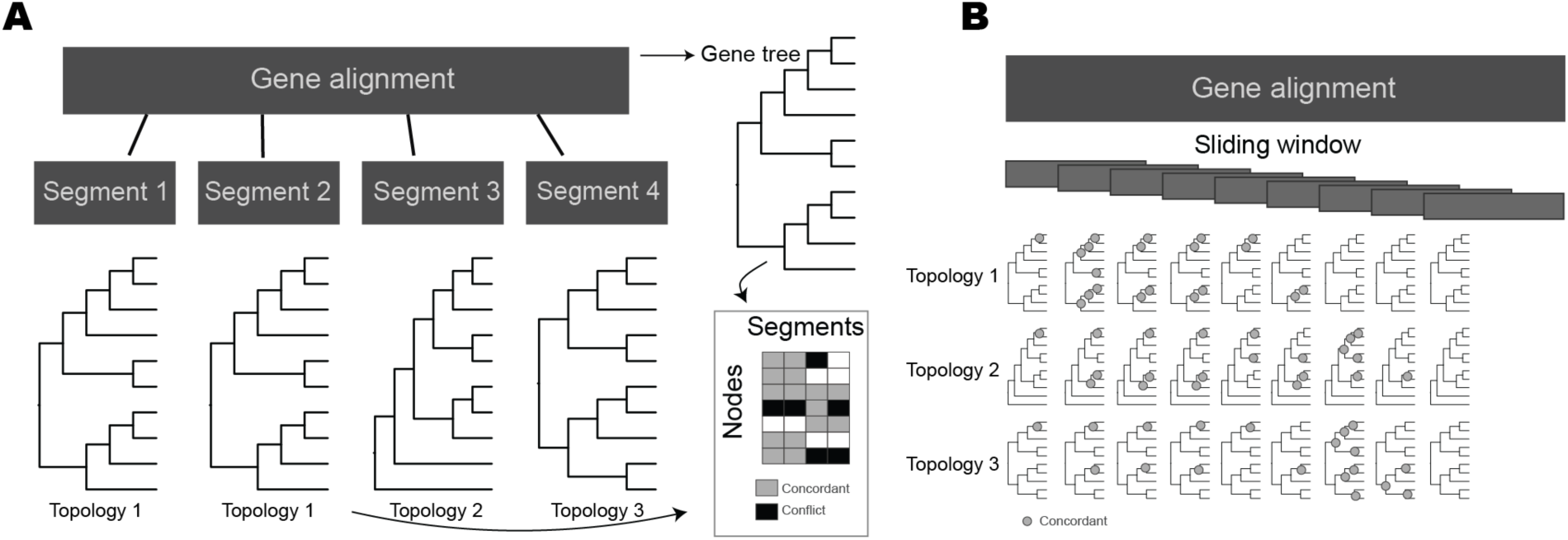
A) Analysis of gene segments and comparisons between gene segment trees and the gene tree recorded in the table with white boxes showing no support, grey showing support, and black showing conflict. These are non-overlapping segments. There are three unique topologies found with Segment 1 and 2 displaying the same topology. B) Sliding window analysis for a more detailed look at where clades display well supported conflict between the gene tree and the segment trees. Support for clades in the three unique topologies are displayed across the sliding window lengths. The sliding window has overlapping portions.

### Simulations

We tested the sensitivity of the approach described above for type I error rate under a variety of scenarios and for positive identification of conflict when present. To do this, we simulated several alignments and analyzed them using our procedure. Each simulation was replicated 100 times.

First, to test for false positive identification under no conflict, we simulated 25-taxon trees using pxbdsim with a birth rate of 1 and a death rate of 0. We then scaled the tree to a root height of 0.75 and simulated 3000 bp under JC with INDELible (Fletcher and Yang, 2009) on the same tree. We conducted a similar simulation but with 50-taxon trees and a root height of 1, and with 25 taxa and 1500 bp. To increase model complexity and test for the impact of differing molecular models, we conducted a simulation with 25 taxa, 3000 bp and different GTR models for first 2000 bp and final 1000 bp (0.6, 0.4, 0.2, 0.8, 1.2 with state freqs 0.3, 0.4, 0.1, 0.2 vs. 0.2, 0.4, 0.6, 0.8, 1.2 with state freqs 0.1, 0.2, 0.3, 0.4), simulated on the same tree. Finally, to generalize this we conducted a simulation with 25 taxa, 3000 bp and different, randomly parameterized GTR models for each 1000 bp segment. We generated GTR parameters by random draws from an exponential distribution with scale = 1, with base frequencies drawn randomly from a uniform distribution and scaled to sum to 1.

For methods which use information criterion to determine the existence and location of breakpoints (Kosakovsky Pond et al. 2006a, b), sectional rate shifts may mislead inference because fit will be improved by separate models which nonetheless correspond to the same topology. We therefore conducted simulations to test the sensitivity of our approach to sectional rate shifts. We generated 25-taxon trees and 3000 bp in 1000 bp segments, where each segment received a different randomly parameterized GTR model as above. Each segment was simulated on the same tree, but scaled such that the root height was 0.5, 0.75 and 1.0, respectively.

We reasoned that alignment error could induce false positives, so in addition to testing our method on simulated alignments, we also simulated under a variety of indel parameters and realigned using MAFFT with defaults (Katoh and Standley, 2013), FSA with defaults (Bradley et al. 2009) and PRANK (-iterate=5) (Löytynoja and Goldman, 2008). We conducted 3 sets of simulations using a negative binomial indel model with r = 1 and p = 0.25 and differing insertion and deletion rates. In each case, we generated a 25-taxon tree as above, and then simulated 3 1000 bp segments under a randomly parameterized GTR model, alongside the specified indel model. In the first set of simulations, the insertion and deletion rate were equal, with 0.03, 0.01, and 0.005 respectively. In the second two sets of simulations, the insertion and deletion rate differed. In the first, the first segment had insertion = 0.03 and deletion = 0.04, the second 0.02 and 0.01 and the third 0.003 and 0.006. In the second, the first segment had insertion = 0.04 and deletion = 0.03, the second 0.01 and 0.02 and the third 0.006 and 0.003.

We conducted simulations with conflict to test the efficacy of our method. In each, we simulated alignments under a single randomly parameterized GTR, but a separate conflicting 25-taxon tree for one segment. We conducted four simulations. In the first we simulated 3000 bp where the last 200 bp were simulated under a conflicting topology. In the second we simulated 3000 bp where the last 500 bp were simulated under a conflicting topology. In the third, we simulated 3000 bp where the last 1000 bp were simulated under a conflicting topology. Finally, we simulated 1500 bp where the last 200 bp were simulated under a conflicting topology.

### Specific examples

To examine the patterns of conflict within genes in more detail, we identified four examples where the conflict identified was not a result of obvious errors (i.e. the examples are not representative of all the inferred conflicts). For each gene examined in more detail we conducted several additional analyses. First, we calculated site-specific *lnL* for each tree constructed from the 1000bp segments, called the segment trees. These values were then compared to site-specific *lnL* of the maximum-likelihood tree constructed from the entire dataset. The site-specific *lnL* calculates the likelihood each site has for a maximum likelihood (ML) topology (Castoe et al. 2009), the likelihood of each site can then be compared between multiple topologies to identify the degree of which the likelihood supports one topology over another (Delta SS *lnL*). By performing this analysis, we were able to examine the degree to which each site supports the segment trees vs. the ML tree. This allowed us to both quantify the degree of significance a site has for an ML tree and whether the regions of genes show bias in their support across topologies. We also conducted sliding-window analyses summing the site-specific *lnL* of the segment trees for 100 bp windows every 20 bp (Fig. 1B). This analysis was performed to help determine the significance of the conflict. For each window we calculated the difference between the maximum and the minimum *lnL* and if the difference was < 0.05, we considered the window to be uninformative. Otherwise, the segment trees that were within 2 *lnL* of the maximum *lnL* were recorded for the window and concordant edges were summarized and reported for each window. We considered these edges to be supported by the window. Finally, we also calculated sliding-window analyses of the base composition to determine if there were any biases in the alignments.

To compare our analyses to previously published methods, we also assessed these specific examples with GARD (Kosakovsky Pond et al., 2006a, b) and phyML_multi (Bossau et al., 2009). For GARD, we used GTR+ Γ and repeated each run in triplicate. Because the outgroup in the respective studies were rarely monophyletic for inferred segment trees, we arbitrarily selected one member of the outgroup to root each tree. Most runs were conducted in HYPHY v2.5.5, but some could not complete due to errors in likelihood calculation, and these were instead conducted in HYPHY v2.5.8. For phyML_multi, we analyzed each gene using TN93+ Γ, because GTR + Γ was not available. Following our segment trees (above), we optimized three trees for EOG2711, 2798 and ENSG00000074803, and two trees for 131. We ran phyML_multi using both the mixed model and HMM approach, but due to issues with the python library used to process results only mixed model results are presented here. To further explore the possible impact of alignment error in these specific examples, we realigned each gene with FSA (--nucprot) and PRANK (-iterate=5 -translate) and analyzed with GARD and phyML_multi as above. For phyML_multi we optimized two or three trees as above for comparative purposes, even if the alignment was longer. Analyses using the detailed method (in Fig 2) failed due to the significant addition of gaps in the altered alignment.

**Figure 2.**
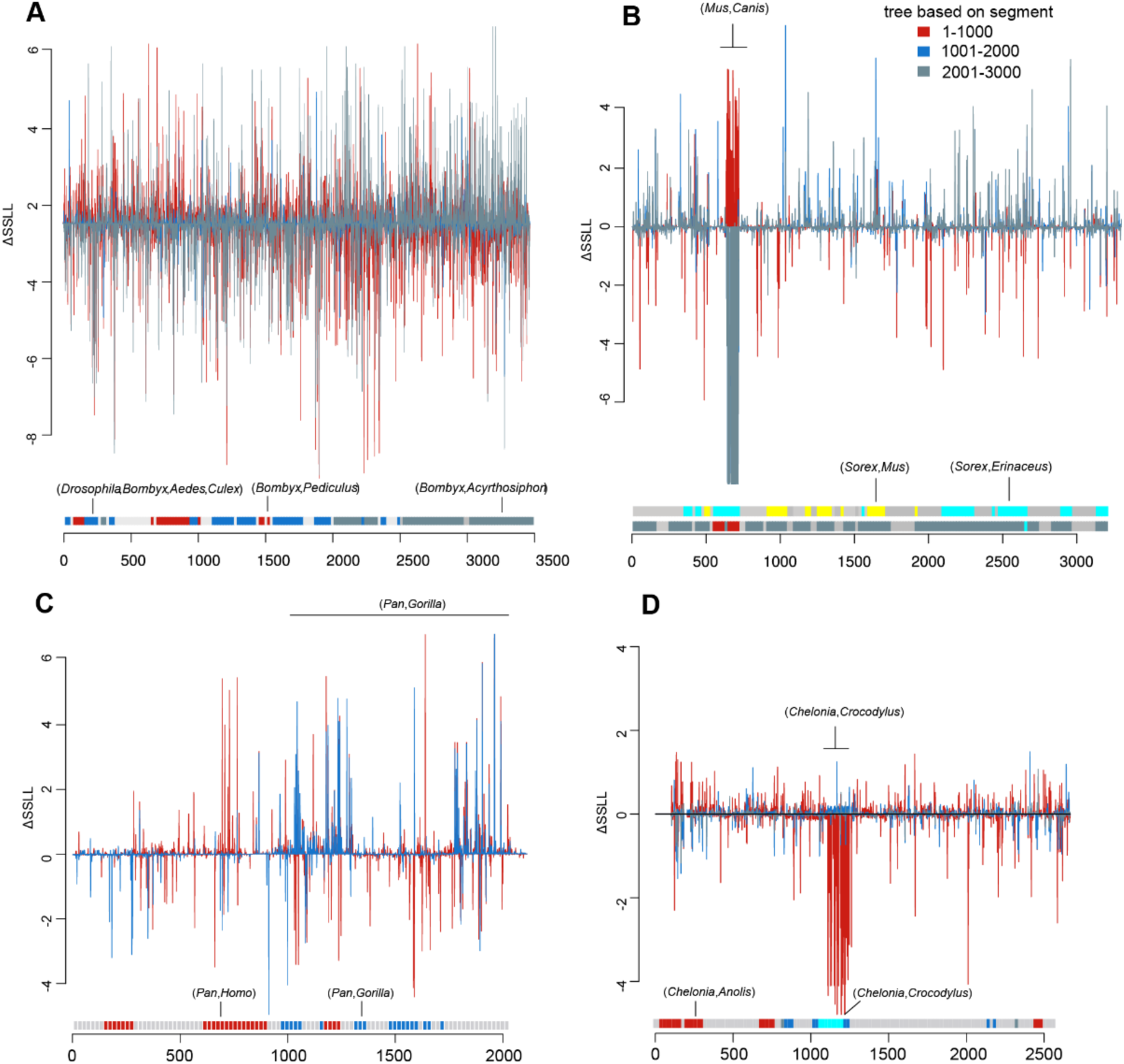
Detailed analysis of within-gene conflict. Delta site-specific *lnL* (ΔSSLL) for each site and the set of segment trees estimated for each dataset are show. Colors/shades represent the ΔSSLL for the tree estimated from the denoted segments. Major deviations for contiguous segments are denoted in B,C,D. Below each plot are the results for the sliding window analysis (using 100 bp segments each 20 bp) showing support for clades and how they change throughout the alignment. Light grey denotes that there was no strong support for any clade in that section. A) *EOG2711* gene from the BUGS dataset. B) *ENSG00000074803* gene in the MAM1 dataset C) *131* gene in the MAM2 dataset D) *2798* gene in the VERT dataset.

## Results

### Simulation Results

Simulations results are presented in Table 1. Our approach has little to no false positives in simple simulations where no conflict is present within genes. This included the situation where data were generated under two different GTR models, or under three randomly parameterized GTR models. Our approach also produced no false positives in the presence of rate shifts on the same tree within a gene. Incorporating indels in the simulation process also led to few false positives. However, realignment of sequences simulated under indel models increased false positive rates. MAFFT and PRANK realignments showed consistently more false positive results with FSA realignments showing fewer false positives. Simulations with conflict indicated that our method was sensitive in cases where conflicting signal made up a large proportion of the overall alignment, with 98% true positives in the case where 1000 bp were generated under a conflicting tree in a 3000 bp alignment. Our method was less sensitive when conflicting signal made up a smaller proportion of the alignment, with only 19.5% true positives in the case where 200 bp were generated under a conflicting tree in a 3000 bp alignment, but 51% true positives in the case where 200 bp were generated under conflicting tree in a 1500 bp alignment. Generally, this demonstrated that this method was effective in identifying conflict when present and with a very low false positive rate.

### Empirical Datasets

We analyzed thirteen datasets for intragenic conflict and found variable results regarding the proportion of those genes with conflict (Table 2). We noted that as taxon sampling increased, so did the inferred conflict. This is, in part, expected because as the number of taxa increases, complexity due to potential errors or biology may be assumed to increase. Nevertheless, this should be explored further. Several datasets consisted of genes that could not be analyzed due to short alignment length.

We analyzed four genes in more detail to better document patterns that are not the result of obvious data assembly errors.

### Invertebrate example

We analyzed the EOG2711 gene in the BUGS dataset (Fig. 2A). This gene did not exhibit the pattern using the more conservative tests presented in Table 2. While several relationships disagree between the segment trees and the ML tree, the primary difference involves the placement of *Bombyx* and relatives. In the ML tree, *Bombyx* is placed in a clade with *Aedes, Culex*, and *Drosophila*. This is consistent with the second tree segment but conflicts with the first and third that have *Bombyx* sister to *Pediculus* and sister to *Acyrthosiphon* respectively GARD inferred between nine and 15 breakpoints. The different segment trees represent a diversity of topologies, with several conflicting in the placement of *Bombus* and *Acyrthosiphon* among others (**Figs. S1-3**). For all runs, the model allowing separate trees had lower AICc than that allowing separate partitions. phyML_multi had optimal support for three trees over 86 segments (**Fig. S4**). Most segments supported a tree with a placement of *Bombyx* more like the ML tree, while the other two trees differed in the placement of *Harpegnathos* and *Trilobium*, among others (**Fig. S4**).

### Mammal examples

We analyzed the ENSG00000074803 gene in the MAM1 dataset (Fig. 2B). The primary conflicts involved the placement of *Mus, Canis, Sorex*, and *Erinaceus*. The ML tree for the entire gene placed *Mus* sister to a clade of seven taxa, *Sorex* sister to *Erinaceus*, and *Canis* sister to *Ailuropda* and *Mustela*. However, the first 1000 bp place *Mus* sister to *Canis* and the second 1000bp place *Sorex* sister to *Mus*. The striking pattern of support for *Mus* sister to *Canis* was not accompanied by a notable alignment or homology difference, as measured by BLAST analyses using the nr dataset. Over three runs, GARD inferred between five and eight breakpoints, with a diversity of topologies (**Figs. S5-S7**). Several conflicted in the placement of *Mus, Sorex, Erinaceus* and *Canis*, among others. For all runs, the model allowing separate trees had far lower AICc than that allowing separate partitions only. phyML_multi had optimal support for only two of three trees over three segments, with the majority supporting a tree which placed *Erinaceus* sister to *Sorex* and *Mus* sister to *Echinops* (**Fig. S8**). A small number of sites supported a tree placing *Mus* sister to *Canis*, and *Erinaceus* sister to *Echinops* (**Fig. S8**).

We analyzed the 131 gene in the MAM2 dataset (Fig. 2C). This gene was not flagged by the conservative tested noted above and reflects an instance where detailed analyses can identify more subtle intragenic conflict. The second segment conflicted with the ML tree in the placement of *Pan* with *Gorilla* instead of *Pan* with *Homo*. Triplicate GARD runs determined support for three trees in only one analysis, with one placing *Gorilla* similarly to the ML tree, and another placing it close to *Echinops* and *Dasypus*, among other conflicts (**Fig. S9**). The model allowing separate trees had far lower AICc than that allowing separate partitions only. phyML_multi optimised support for two trees over 28 segments (**Fig. S10**). These trees similarly conflicted with one another and the ML tree in primate relationships, with the first placing *Pan* sister to *Homo* as in the ML tree, and the second placing *Pan, Gorilla* and *Homo* in a grade leading to *Callithrix* and *Pongo* among other conflicts.

### Vertebrate example

We analyzed the 2798 gene in the VERT dataset (Fig. 2D). The primary conflicts between the trees based on 1000bp segments and the ML analyses involved the placement of crocodile and sea turtle. The ML analyses supported sea turtle as sister to soft turtle. However, the second 1000bp segment supported sea turtle as sister to crocodile. The dramatic site likelihood shift at ∼1100-1300 bp (Fig. 2D) was not associated with any notable homology or alignment problem. GARD inferred between 17 and 19 segments with topologies varying in the placement of sea turtle, soft turtle, and *Anolis* (**Figs. S11-S13**). For all runs, the model allowing separate trees had far lower AICc than that allowing separate partitions. phyML_multi had optimal support for three trees over 10 segments, with the majority of sites supporting one of two trees placing sea turtle sister to *Anolis*, and a small number of sites supporting sea turtle sister to crocodile (**Fig. S14**).

For all exemplars, analysis of realignments continued to exhibit breakpoints featuring similar conflicts with the ML tree, albeit at different positions (**Fig. S15-45**).

## Discussion

We have demonstrated that intragenic conflict can be common in empirical datasets. Some of the datasets analyzed here overlap in taxon sampling but vary greatly in the frequency of intragenic conflict (e.g., MAM1 and MAM2) suggesting that, while biological processes may play a role in generating conflict (Mendes et al. 2019), non-biological errors are likely to be a major source of incongruence. For example, MAM2 has been thoroughly analyzed to uncover errors exist within the dataset assembly (Springer and Gatesy, 2018). Importantly, all the phylogenetic methods from the original publications for each dataset in Table 2 assumed a single topology underlying the alignment. Our analyses suggest, that for many gene regions this would represent model misspecification, as there are several well supported topologies underlying many genes which may mislead analyses on these data. Several of the taxon-rich datasets, including FISH and FROG, had very high levels intragenic conflict across the genes we analyzed. This points to either systematic errors in dataset assembly, extensive recombination, or other biological processes. We note that, while there was a tendency for taxon-rich datasets to exhibit these issues, the PLAN dataset did not exhibit a particularly high rate of conflicts. Nevertheless, we expect that increasing taxon sampling may potentially result in more intragenic conflict for biological or non-biological reasons as increasing sampling will necessarily increase biological heterogeneity (for example the probability of sampling a recombination event) and the potential for error. Additionally, longer alignments would be expected to harbor more recombination breakpoints. Our analyses filtered datasets for regions that could produce at least two gene segments and so we could potentially bias upward our estimates of within-gene conflict. However, we did not notice a trend of datasets with longer gene regions to exhibit more conflict overall, and much smaller genes are expected to yield poor-quality phylogenetic inference. Nonetheless, the addition of both more taxa and more sites will increase the potential for biological and systematic within-gene conflict.

Non-biological errors may arise from alignment inaccuracies, homology issues, and errors in dataset assembly perhaps exacerbated for large datasets assembled using significant automation. However, while errors may be a major source of intragenic conflict, the data examined in Fig. 2 did not in general exhibit obvious errors (2798 from the VERT dataset could perhaps plausibly include misannotated sequences, but it is difficult to confirm this without a more in-depth examination of the soft-shell turtle genome), and breakpoints were detected with multiple methods even after re-alignment with different algorithms. It is still unclear what the source of the conflict is in those specific examples. However the conflict we detect, without obvious systematic error, contributes to a growing body of literature that is discovering intragenic conflict due to biological processes such as recombination (e.g., Scornavacca and Galtier 2016, Mendes et al. 2019). Without having generated the original datasets, including assembly or alignment, it would be very difficult to untangle what the source of conflict was in each case. However, in order to ensure that biological conclusions were not the result of noise and error, it would be important to determine whether intragenic heterogeneity was being properly accounted for in these empirical datasets.

The importance of accurate alignment for tree inference is well-understood (Ogden and Rosenberg, 2006). The results we present here suggest that alignment error impacts not only maximum likelihood topology estimates but can also induce intragenic conflict. Specifically, in our simulations, realignment of simulated data containing indels led to increased false positives. Realignment with a progressive algorithm (MAFFT), which is known to over-align (Katoh et al. 2016, Vialle et al. 2018), produced greater numbers of false positives. By contrast FSA, which typically under-aligns (Bradley et al. 2009), led to lower rates of false positives. This inference is supported by analysis of empirical datasets. For example, the CARN dataset from Walker et al. (2017) was aligned with PRANK and showed comparatively lower rates of within-gene conflict. Conversely, although realignment with more accurate aligners reduced false positive occurrence in simulations, multiple trees were still inferred for exemplars when realigning, suggesting true signals of multiple trees that are not due to alignment errors. Despite the overall simplicity of our simulation process, our rate shift experiments and changes in indel rates between segments should mimic some of the more complex dynamics occurring in e.g. genes encoding multi-domain proteins.

Whether due to biological processes or errors in dataset assembly, intragenic conflict should not be ignored. Intragenic conflict can drive false positives in positive selection analyses (Anisimova et al., 2003), influence branch length estimation, and bias phylogenetic reconstruction. These and other errors are to be expected as intragenic conflict violates typical phylogenetic models that assume a single tree for the length of the alignment. Some have suggested that recombination may have a minimal impact on species tree inference (Lanier and Knowles, 2012). However, others have noted that only a few conflicting sites can drive tree inference (Shen et al. 2017). Furthermore, our results have implications for analyses that assume that a gene region, regardless of data type, is the most meaningful phylogenetic unit (e.g., summary coalescent analyses). In the presence of intragenic conflict, an entire gene may not be a meaningful unit for phylogenetic analysis.

Our method presents one way to highlight general problems with dataset assembly and to identify important molecular evolutionary patterns. Importantly, the methods presented here do not assume a particular source of conflict. Nevertheless, if a dataset exhibits significant intragenic conflict, it warrants further investigation regardless of the source. Biological sources of intragenic conflict can provide important information about molecular evolutionary processes, but in order to better understand these processes we need to ensure that we identify biological conflict and not error. Disentangling the sources of intragenic conflict will lead to cleaner datasets, more robust species tree inferences, and a greater understanding of the molecular evolutionary processes shaping genomes..

## Acknowledgments

We would like to thank James Pease, Jeremy Beaulieu, Jonathan Chang, and Caroline Parins-Fukuchi for helpful comments. NWH was supported by a Woolf Fisher Cambridge Scholarship. JFW was supported by University of Michigan Rackham Pre-doctoral fellowship. SAS was supported by a University of Michigan MICDE pilot grant and NSF DEB 1917146.

## References

Allman ES, Kubatko LS, Rhodes JA. 2017. Split Scores: A Tool to Quantify Phylogenetic Signal in Genome-Scale Data. Systematic Biology. 66(4):620–636. doi: 10.1093/sysbio/syw103.

Ané C. 2011. Detecting Phylogenetic Breakpoints and Discordance from Genome-Wide Alignments for Species Tree Reconstruction. Genome Biology and Evolution. 3:246–258. doi: 10.1093/gbe/evr013.

Anisimova M, Nielsen R, Yang Z. 2003. Effect of Recombination on the Accuracy of the Likelihood Method for Detecting Positive Selection at Amino Acid Sites. Genetics. 164(3):1229.

Boussau B, Guéguen L, Gouy M. 2009. A mixture model and a hidden markov model to simultaneously detect recombination breakpoints and reconstruct phylogenies. Evol Bioinform Online. 5:67–79.

Boussau B, Szöllősi GJ, Duret L, Gouy M, Tannier E, Daubin V. 2013. Genome-scale coestimation of species and gene trees. Genome Research. 23(2):323–330. doi: 10.1101/gr.141978.112.

Bradley RK, Roberts A, Smoot M, Juvekar S, Do J, Dewey C, Holmes I, Pachter L. 2009. Fast statistical alignment. PLoS Comput Biol. 5(5):e1000392. doi: 10.1371/journal.pcbi.1000392.

Brown JM, Thomson RC. 2016. Bayes Factors Unmask Highly Variable Information Content, Bias, and Extreme Influence in Phylogenomic Analyses. Systematic Biology. 66(4):517–530. doi: 10.1093/sysbio/syw101.

Brown JW, Walker JF, Smith SA. 2017. Phyx: phylogenetic tools for unix. Bioinformatics. 33(12):1886–1888. doi: 10.1093/bioinformatics/btx063.

Castoe TA, de Koning APJ, Kim H-M, Gu W, Noonan BP, Naylor G, Jiang ZJ, Parkinson CL, Pollock DD, Hillis DM. 2009. Evidence for an Ancient Adaptive Episode of Convergent Molecular Evolution. Proceedings of the National Academy of Sciences of the United States of America. 106(22):8986–8991.

Chen M-Y, Liang D, Zhang P. 2017. Phylogenomic Resolution of the Phylogeny of Laurasiatherian Mammals: Exploring Phylogenetic Signals within Coding and Noncoding Sequences. Genome Biology and Evolution. 9(8):1998–2012. doi: 10.1093/gbe/evx147.

Edwards SV. 2009. Is a New and General Theory of Molecular Systematics Emerging? Evolution. 63(1):1–19. doi: 10.1111/j.1558-5646.2008.00549.x.

Feng Y-J, Blackburn DC, Liang D, Hillis DM, Wake DB, Cannatella DC, Zhang P. 2017. Phylogenomics reveals rapid, simultaneous diversification of three major clades of Gondwanan frogs at the Cretaceous– Paleogene boundary. Proc Natl Acad Sci USA. 114(29):E5864. doi: 10.1073/pnas.1704632114.

Fletcher W, Yang Z. 2009. INDELible: A Flexible Simulator of Biological Sequence Evolution. Molecular Biology and Evolution. 26(8):1879–1888. doi: 10.1093/molbev/msp098.

Gatesy J, Springer MS. 2013. Concatenation versus coalescence versus “concatalescence.” Proc Natl Acad Sci USA. 110(13):E1179. doi: 10.1073/pnas.1221121110.

Guindon S, Dufayard J-F, Lefort V, Anisimova M, Hordijk W, Gascuel O. 2010. New Algorithms and Methods to Estimate Maximum-Likelihood Phylogenies: Assessing the Performance of PhyML 3.0. Systematic Biology. 59(3):307–321. doi: 10.1093/sysbio/syq010.

Hoang DT, Chernomor O, von Haeseler A, Minh BQ, Vinh LS. 2018. UFBoot2: Improving the Ultrafast Bootstrap Approximation. Mol Biol Evol. 35(2):518–522. doi: 10.1093/molbev/msx281.

Hobolth A, Christensen OF, Mailund T, Schierup MH. 2007. Genomic Relationships and Speciation Times of Human, Chimpanzee, and Gorilla Inferred from a Coalescent Hidden Markov Model. PLOS Genetics. 3(2):e7. doi: 10.1371/journal.pgen.0030007.

Hughes LC, Ortí G, Huang Y, Sun Y, Baldwin CC, Thompson AW, Arcila D, Betancur-R. R, Li C, Becker L, et al. 2018. Comprehensive phylogeny of ray-finned fishes (Actinopterygii) based on transcriptomic and genomic data. Proc Natl Acad Sci USA. 115(24):6249. doi: 10.1073/pnas.1719358115.

Husmeier D, McGuire G. 2003. Detecting recombination in 4-taxa DNA sequence alignments with Bayesian hidden Markov models and Markov chain Monte Carlo. Mol Biol Evol. 20(3):315–337. doi: 10.1093/molbev/msg039.

Katoh K, Standley DM. 2013. MAFFT Multiple Sequence Alignment Software Version 7: Improvements in Performance and Usability. Mol Biol Evol. 30(4):772–780. doi: 10.1093/molbev/mst010.

Katoh K, Standley DM. 2016. A simple method to control over-alignment in the MAFFT multiple sequence alignment program. Bioinformatics. 32(13):1933–1942. doi: 10.1093/bioinformatics/btw108.

Kosakovsky Pond SL, Posada D, Gravenor MB, Woelk CH, Frost Simon D. W. 2006. Automated Phylogenetic Detection of Recombination Using a Genetic Algorithm. Molecular Biology and Evolution. 23(10):1891–1901. doi: 10.1093/molbev/msl051.

Kosakovsky Pond SL, Posada D, Gravenor MB, Woelk CH, Frost Simon D.W. 2006. GARD: a genetic algorithm for recombination detection. Bioinformatics. 22(24):3096–3098. doi: 10.1093/bioinformatics/btl474.

Lanier HC, Knowles LL. 2012. Is Recombination a Problem for Species-Tree Analyses? Systematic Biology. 61(4):691–701. doi: 10.1093/sysbio/syr128.

Liu Y, Johnson MG, Cox CJ, Medina R, Devos N, Vanderpoorten A, Hedenäs L, Bell NE, Shevock JR, Aguero B, et al. 2019. Resolution of the ordinal phylogeny of mosses using targeted exons from organellar and nuclear genomes. Nature Communications. 10(1):1485. doi: 10.1038/s41467-019-09454-w.

Löytynoja A, Goldman N. 2005. An algorithm for progressive multiple alignment of sequences with insertions. Proc Natl Acad Sci USA. 102(30):10557–10562. doi: 10.1073/pnas.0409137102.

McCormack JE, Faircloth BC, Crawford NG, Gowaty PA, Brumfield RT, Glenn TC. 2012. Ultraconserved elements are novel phylogenomic markers that resolve placental mammal phylogeny when combined with species-tree analysis. Genome Res. 22(4):746–754. doi: 10.1101/gr.125864.111.

Mendes FK, Livera AP, Hahn MW. 2019. The perils of intralocus recombination for inferences of molecular convergence. Philosophical Transactions of the Royal Society B: Biological Sciences. 374(1777):20180244. doi: 10.1098/rstb.2018.0244.

Mirarab S, Reaz R, Bayzid MdS, Zimmermann T, Swenson MS, Warnow T. 2014. ASTRAL: genome-scale coalescent-based species tree estimation. Bioinformatics. 30(17):i541–i548. doi: 10.1093/bioinformatics/btu462.

Nguyen L-T, Schmidt HA, von Haeseler A, Minh BQ. 2015. IQ-TREE: a fast and effective stochastic algorithm for estimating maximum-likelihood phylogenies. Mol Biol Evol. 32(1):268–274. doi: 10.1093/molbev/msu300.

Niehuis O, Hartig G, Grath S, Pohl H, Lehmann J, Tafer H, Donath A, Krauss V, Eisenhardt C, Hertel J, et al. 2012. Genomic and Morphological Evidence Converge to Resolve the Enigma of Strepsiptera. Current Biology. 22(14):1309–1313. doi: 10.1016/j.cub.2012.05.018.

Ogden TH, Rosenberg MS. 2006. Multiple Sequence Alignment Accuracy and Phylogenetic Inference. Syst Biol. 55(2):314–328. doi: 10.1080/10635150500541730.

Pizarro D, Divakar PK, Grewe F, Leavitt SD, Huang J-P, Dal Grande F, Schmitt I, Wedin M, Crespo A, Lumbsch HT. 2018. Phylogenomic analysis of 2556 single-copy protein-coding genes resolves most evolutionary relationships for the major clades in the most diverse group of lichen-forming fungi. Fungal Diversity. 92(1):31–41. doi: 10.1007/s13225-018-0407-7.

Scornavacca C, Galtier N. 2016. Incomplete Lineage Sorting in Mammalian Phylogenomics. Systematic Biology. 66(1):112–120. doi: 10.1093/sysbio/syw082.

Shen X-X, Hittinger CT, Rokas A. 2017. Contentious relationships in phylogenomic studies can be driven by a handful of genes. Nature Ecology & Evolution. 1(5):0126. doi: 10.1038/s41559-017-0126.

Smith SA, Moore MJ, Brown JW, Yang Y. 2015. Analysis of phylogenomic datasets reveals conflict, concordance, and gene duplications with examples from animals and plants. BMC Evolutionary Biology. 15(1):150. doi: 10.1186/s12862-015-0423-0.

Song S, Liu L, Edwards SV, Wu S. 2012. Resolving conflict in eutherian mammal phylogeny using phylogenomics and the multispecies coalescent model. Proc Natl Acad Sci USA. 109(37):14942. doi: 10.1073/pnas.1211733109.

Springer MS, Gatesy J. 2018. On the importance of homology in the age of phylogenomics. Systematics and Biodiversity. 16(3):210–228. doi: 10.1080/14772000.2017.1401016.

Suchard MA, Weiss RE, Dorman KS, Sinsheimer JS. 2002. Oh Brother, Where Art Thou? A Bayes Factor Test for Recombination with Uncertain Heritage. Systematic Biology. 51(5):715–728. doi: 10.1080/10635150290102384.

Vialle RA, Tamuri AU, Goldman N. 2018. Alignment Modulates Ancestral Sequence Reconstruction Accuracy. Mol Biol Evol. 35(7):1783–1797. doi: 10.1093/molbev/msy055.

Walker JF, Brown JW, Smith SA. 2018. Analyzing Contentious Relationships and Outlier Genes in Phylogenomics. Systematic Biology. 67(5):916–924. doi: 10.1093/sysbio/syy043.

Walker JF, Yang Y, Moore MJ, Mikenas J, Timoneda A, Brockington SF, Smith SA. 2017. Widespread paleopolyploidy, gene tree conflict, and recalcitrant relationships among the carnivorous Caryophyllales. American Journal of Botany. 104(6):858–867. doi: 10.3732/ajb.1700083.

Wang Z, Pascual-Anaya J, Zadissa A, Li W, Niimura Y, Huang Z, Li C, White S, Xiong Z, Fang D, et al. 2013. The draft genomes of soft-shell turtle and green sea turtle yield insights into the development and evolution of the turtle-specific body plan. Nature Genetics. 45:701.

Wickett NJ, Mirarab S, Nguyen N, Warnow T, Carpenter E, Matasci N, Ayyampalayam S, Barker MS, Burleigh JG, Gitzendanner MA, et al. 2014. Phylotranscriptomic analysis of the origin and early diversification of land plants. PNAS. 111(45):E4859–E4868. doi: 10.1073/pnas.1323926111.

Wu S, Song S, Liu L, Edwards SV. 2013. Reply to Gatesy and Springer: The multispecies coalescent model can effectively handle recombination and gene tree heterogeneity. Proc Natl Acad Sci USA. 110(13):E1180. doi: 10.1073/pnas.1300129110.

