## Supplemental Information for "Intragenic conflict in phylogenomic datasets"

### Supplementary figures and captions

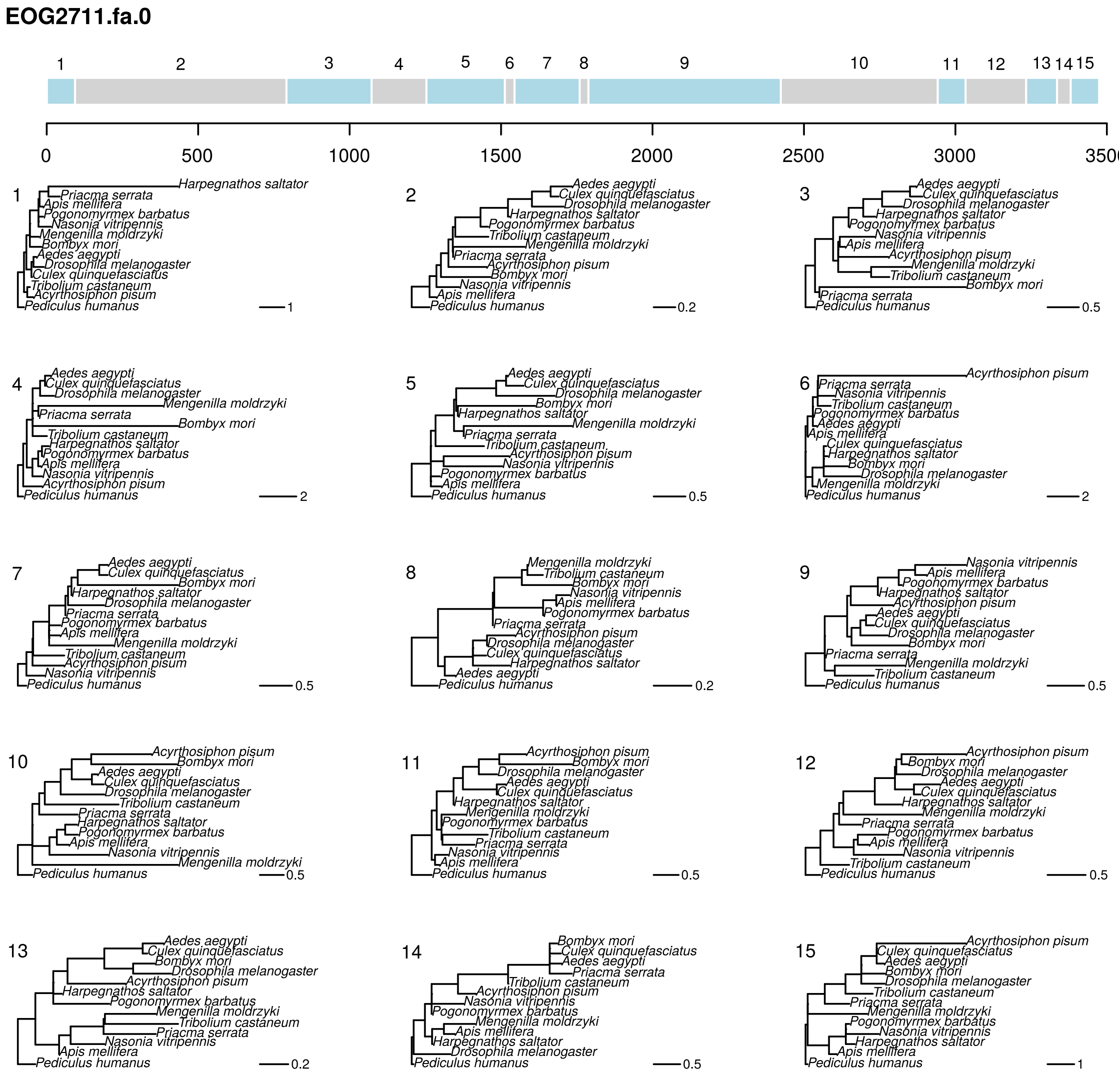


**Figure S1:** replicate one GARD analysis of EOG2711 under GTR+G. Tree topologies correspond to numbered segments, colors are arbitrary. Branch lengths and scale correspond to expected nucleotide substitutions per site.

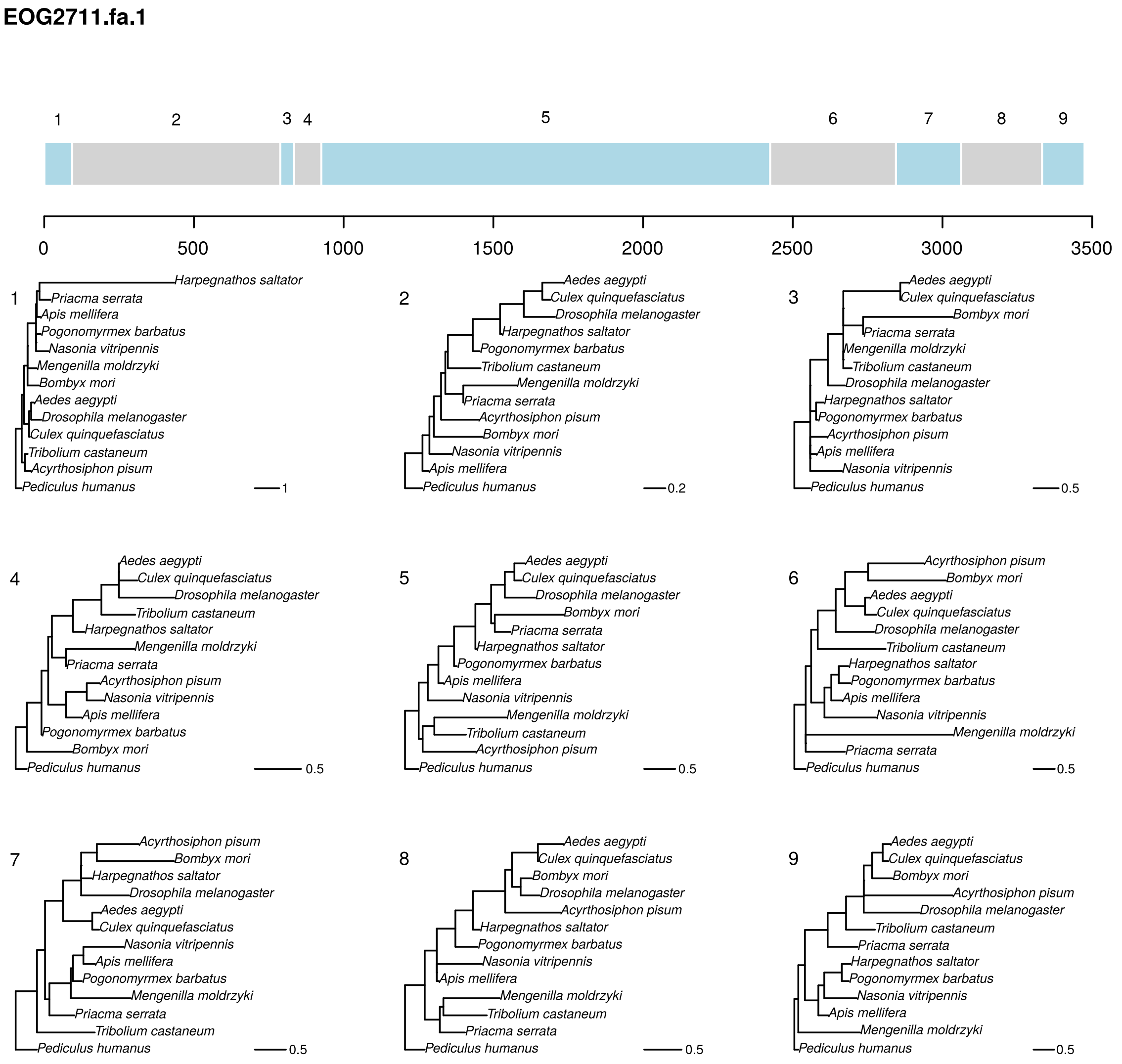


**Figure S2:** replicate two GARD analysis of EOG2711 under GTR+G. Tree topologies correspond to numbered segments, colors are arbitrary. Branch lengths and scale correspond to expected nucleotide substitutions per site.

**EOG2711.fa.2**

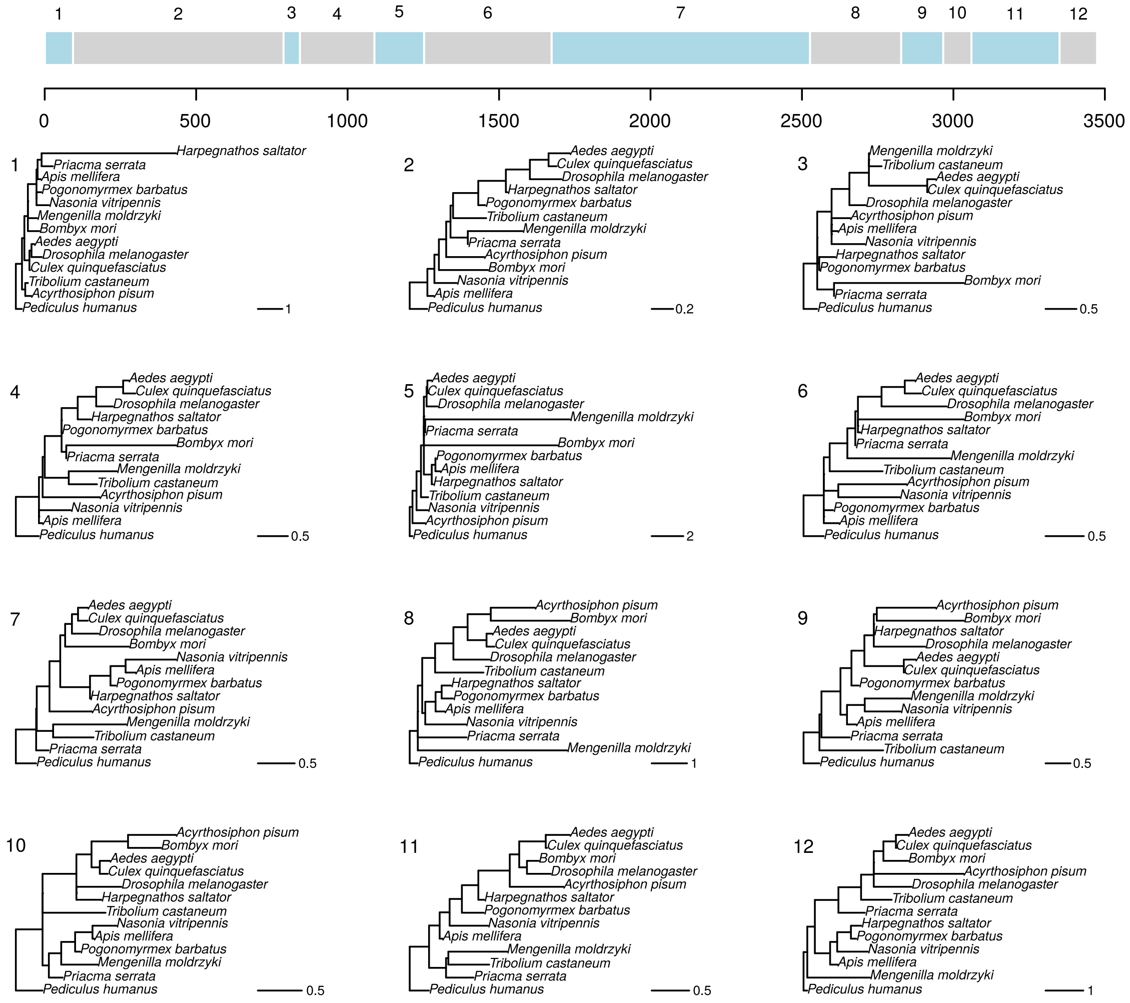


**Figure S3:** replicate three GARD analysis of EOG2711 under GTR+G. Tree topologies correspond to numbered segments, colors are arbitrary. Branch lengths and scale correspond to expected nucleotide substitutions per site.

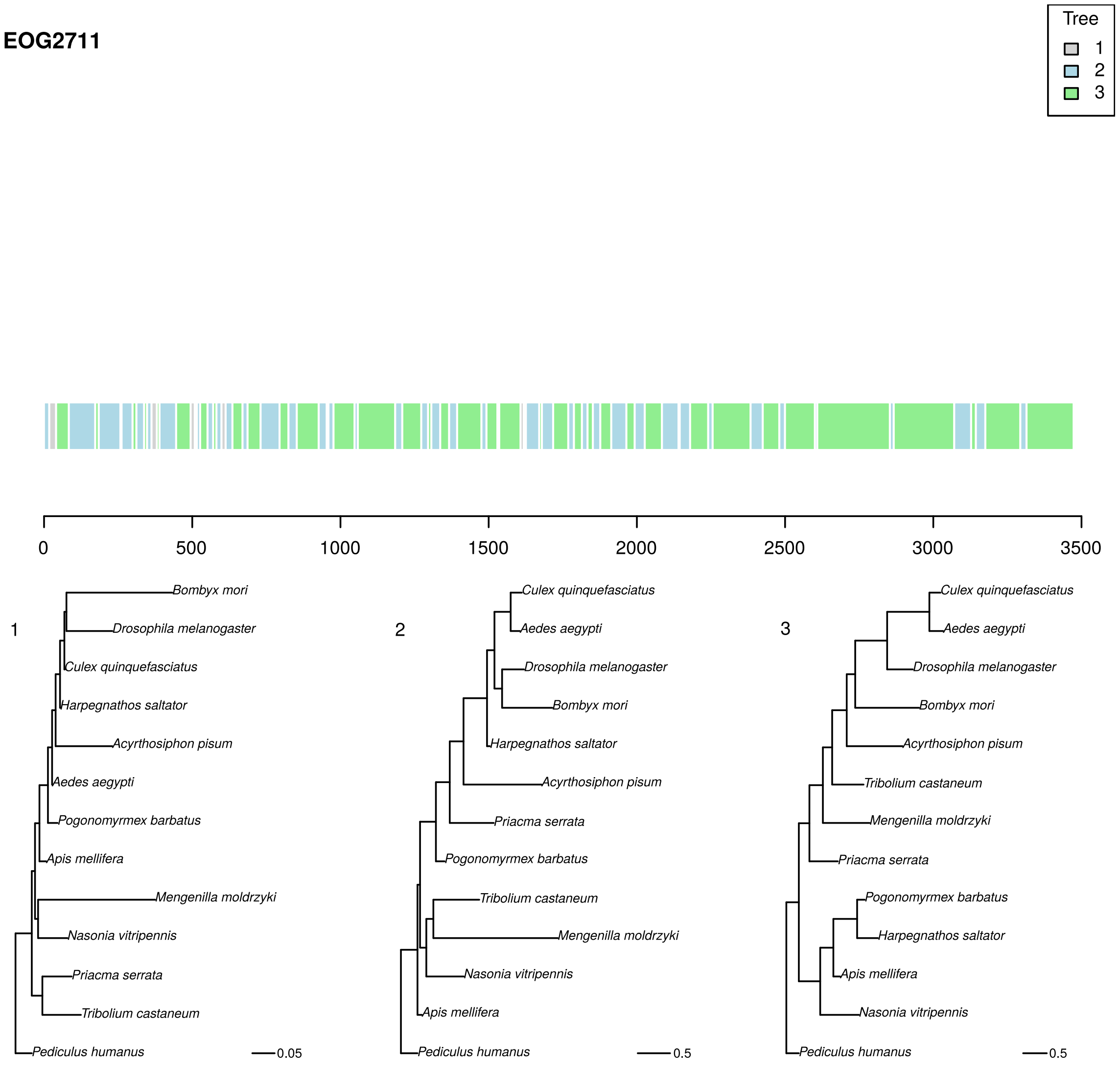


**Figure S4:** phyML_multi analysis of EOG2711 under TN93+G with three trees. Colors correspond to the best-supported tree for each segment. Branch lengths and scale bar correspond to expected nucleotide substitutions per site.

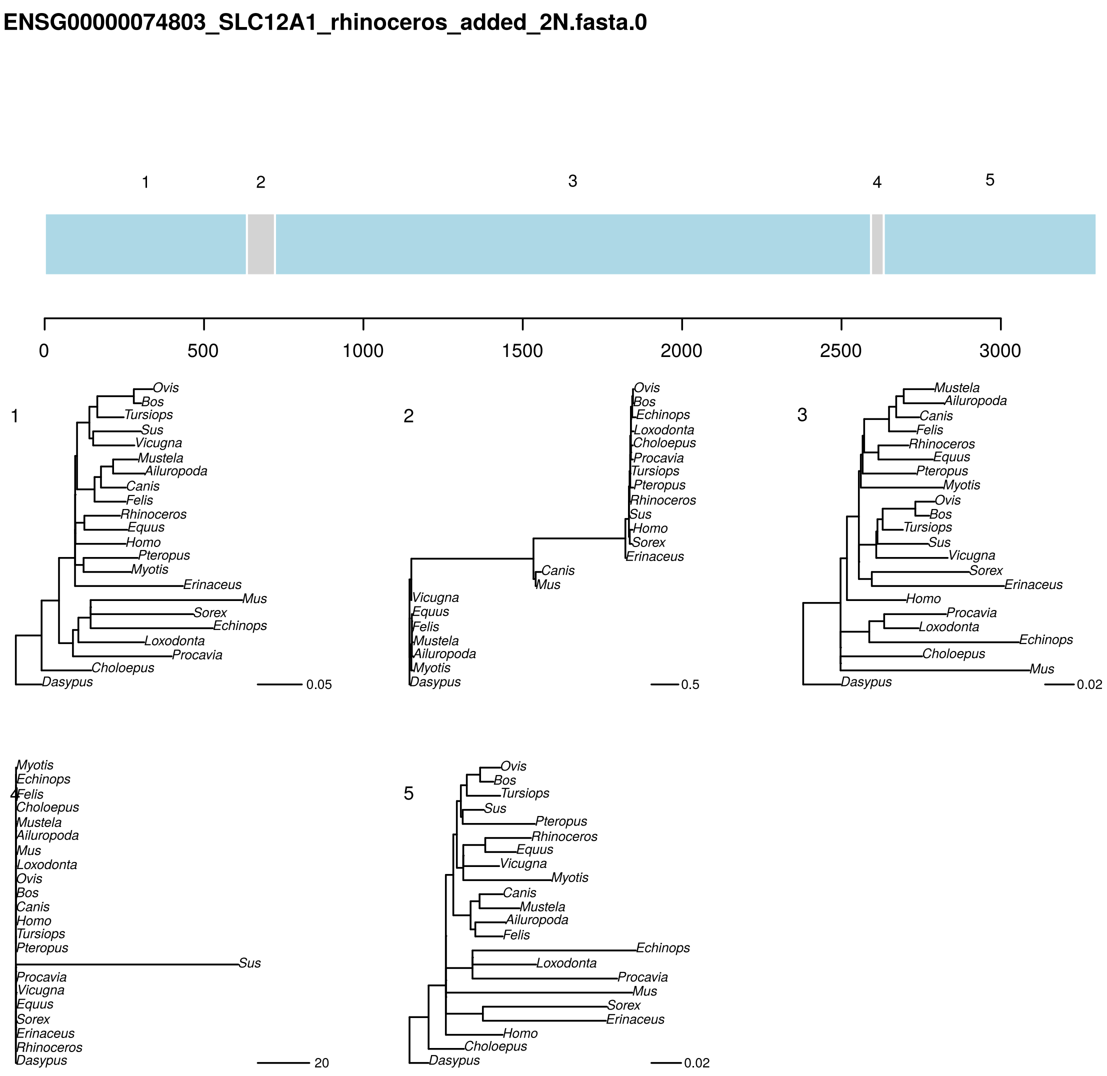


**Figure S5:** replicate one GARD analysis of ENSG00000074803 under GTR+G. Tree topologies correspond to numbered segments, colors are arbitrary. Branch lengths and scale correspond to expected nucleotide substitutions per site.

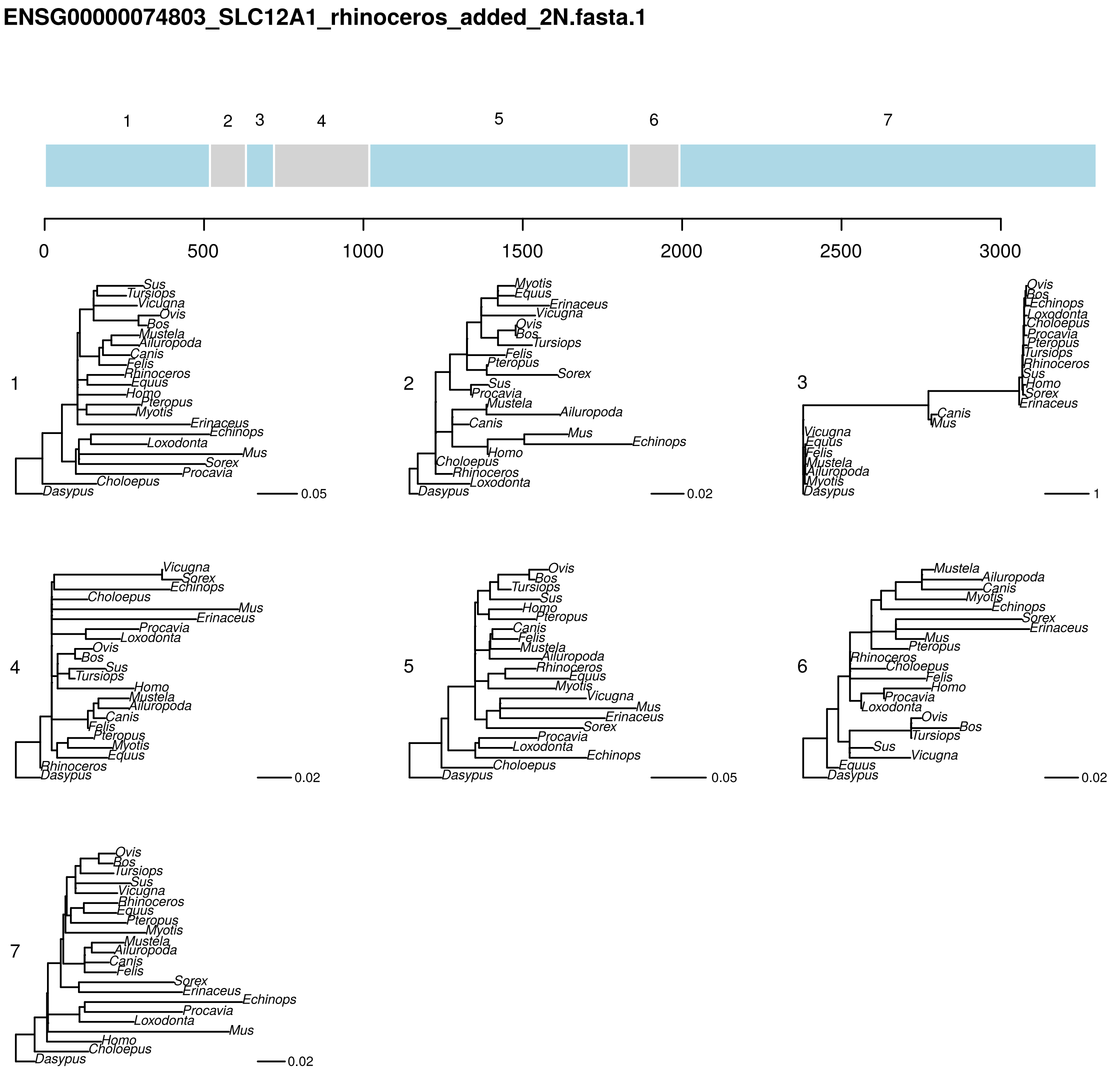


**Figure S6:** replicate two GARD analysis of ENSG00000074803 under GTR+G. Tree topologies correspond to numbered segments, colors are arbitrary. Branch lengths and scale correspond to expected nucleotide substitutions per site.

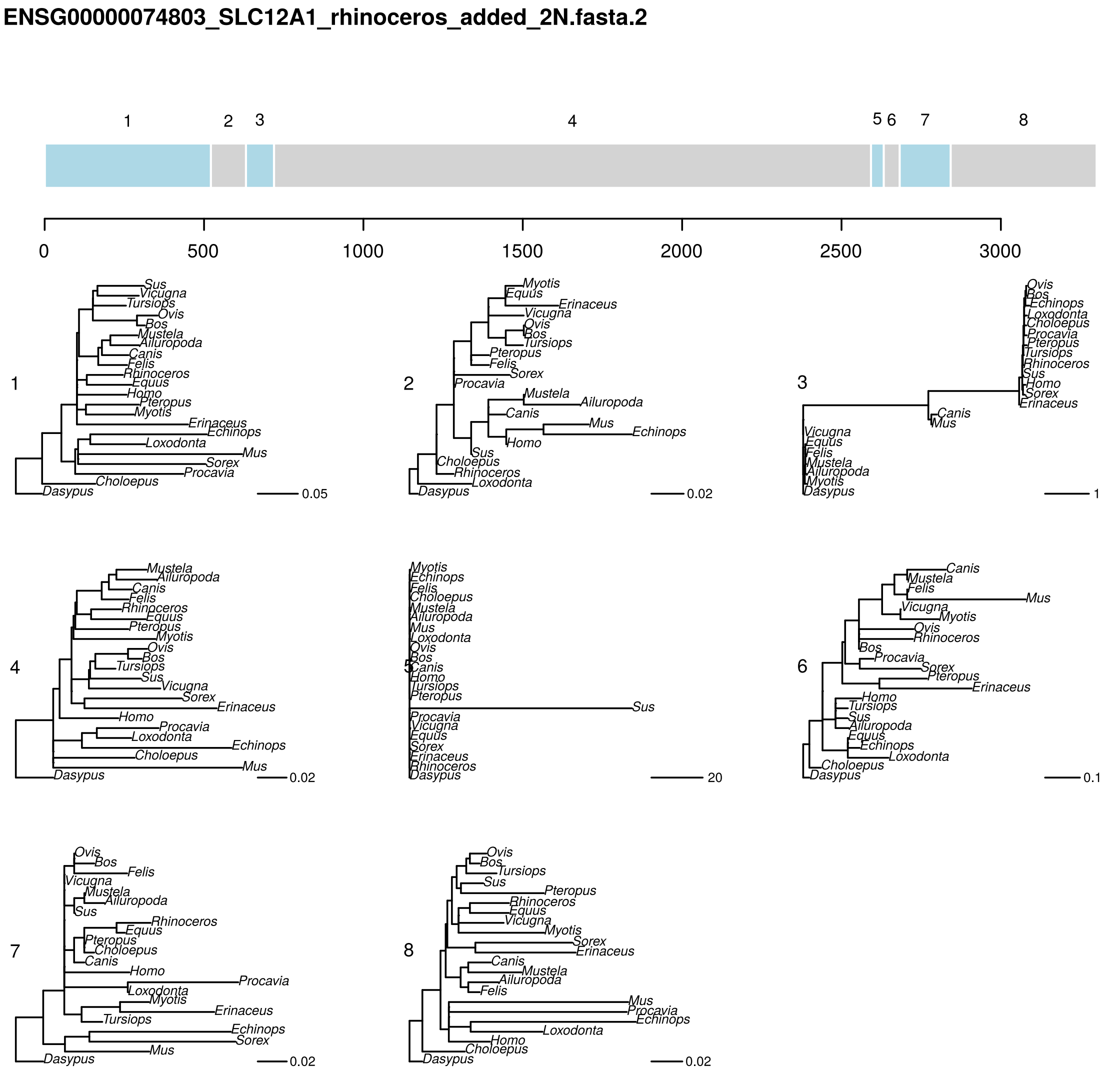


**Figure S7:** replicate three GARD analysis of ENSG00000074803 under GTR+G. Tree topologies correspond to numbered segments, colors are arbitrary. Branch lengths and scale correspond to expected nucleotide substitutions per site.

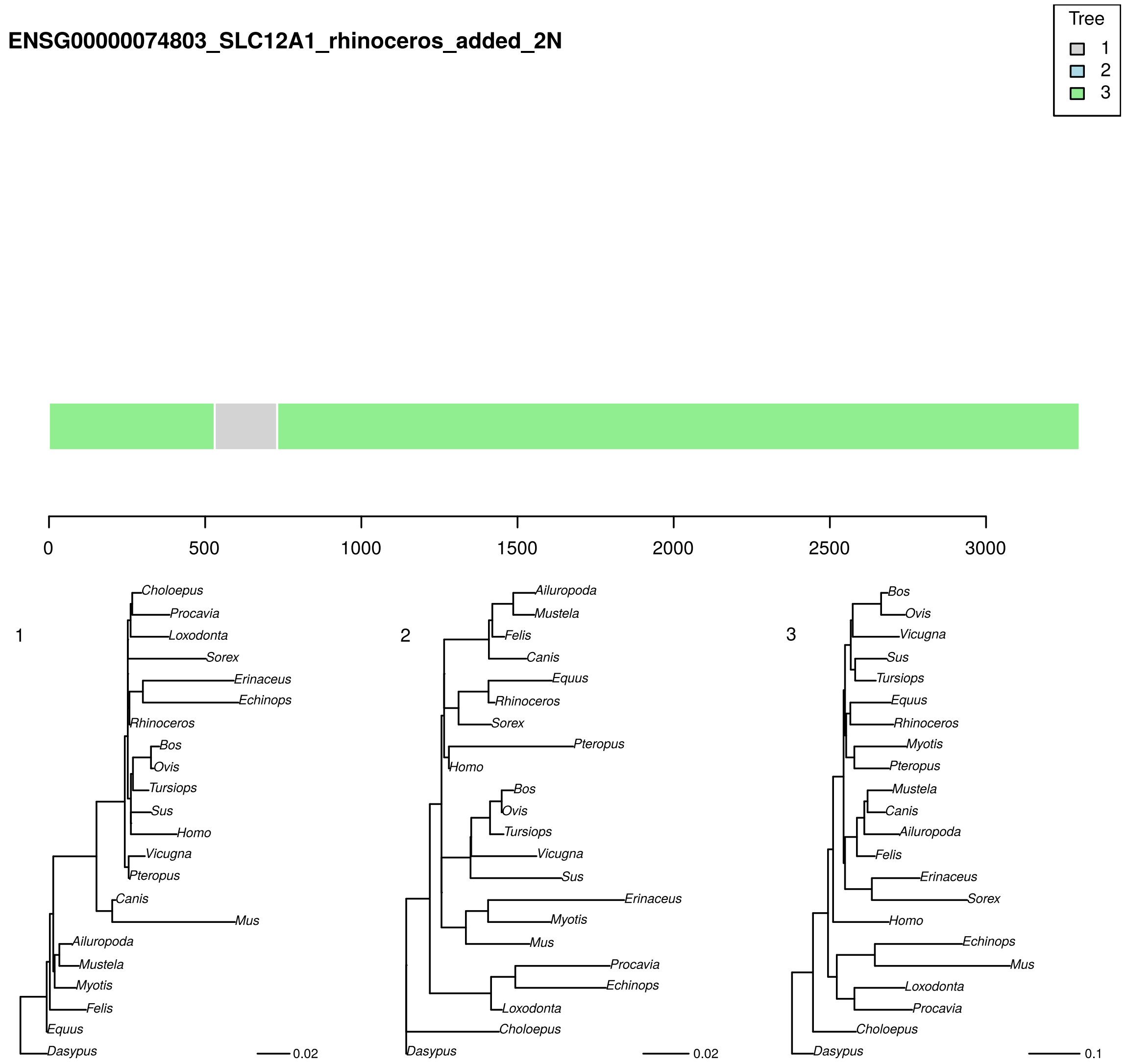


**Figure S8:** phyML_multi analysis of ENSG00000074803 under TN93+G with three trees. Colors correspond to the best-supported tree for each segment. Branch lengths and scale bar correspond to expected nucleotide substitutions per site.

**131.fasta.1**

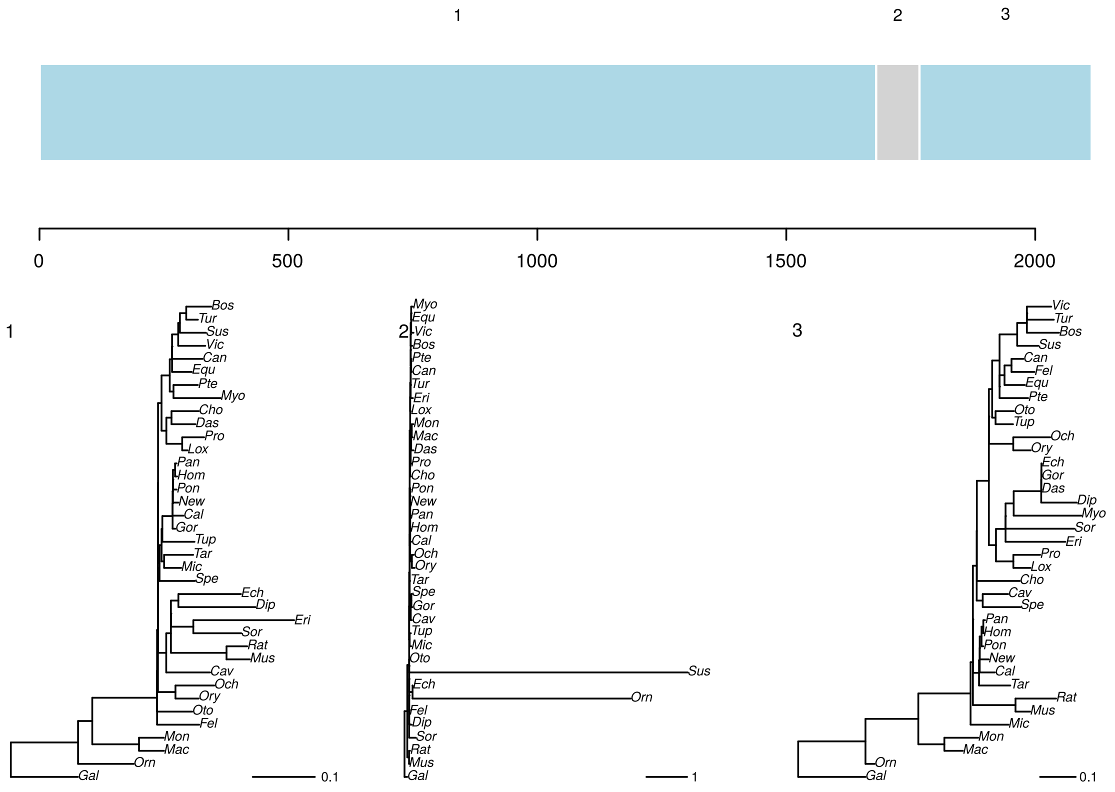

**Figure S9:** GARD analysis of 131 under GTR+G. Tree topologies correspond to numbered segments, colors are arbitrary. Branch lengths and scale bar correspond to expected nucleotide substitutions per site.

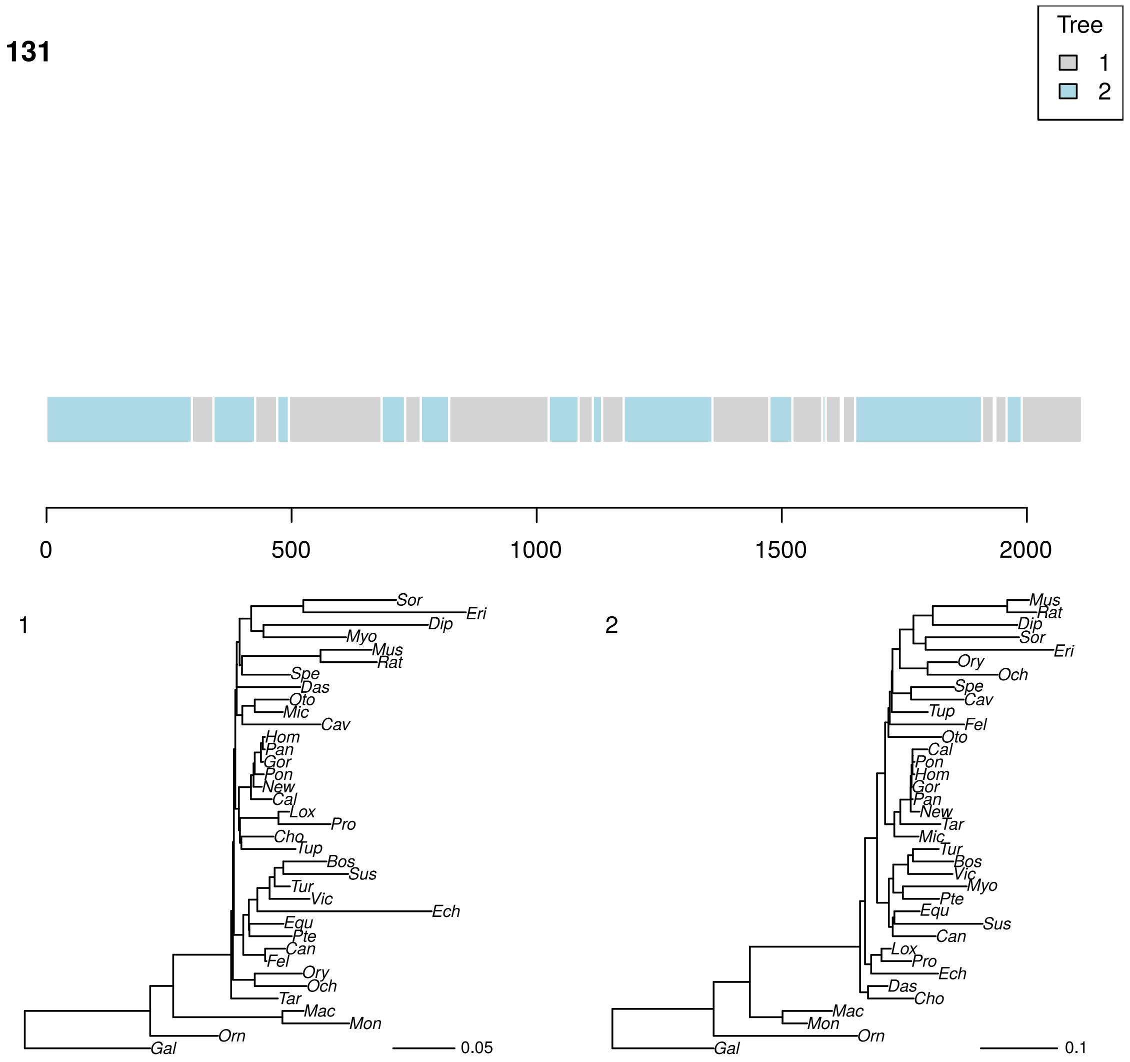


**Figure S10:** phyML_multi analysis of 131 under TN93+G with two trees. Colors correspond to the best-supported tree for each segment. Branch lengths and scale bar correspond to expected nucleotide substitutions per site.

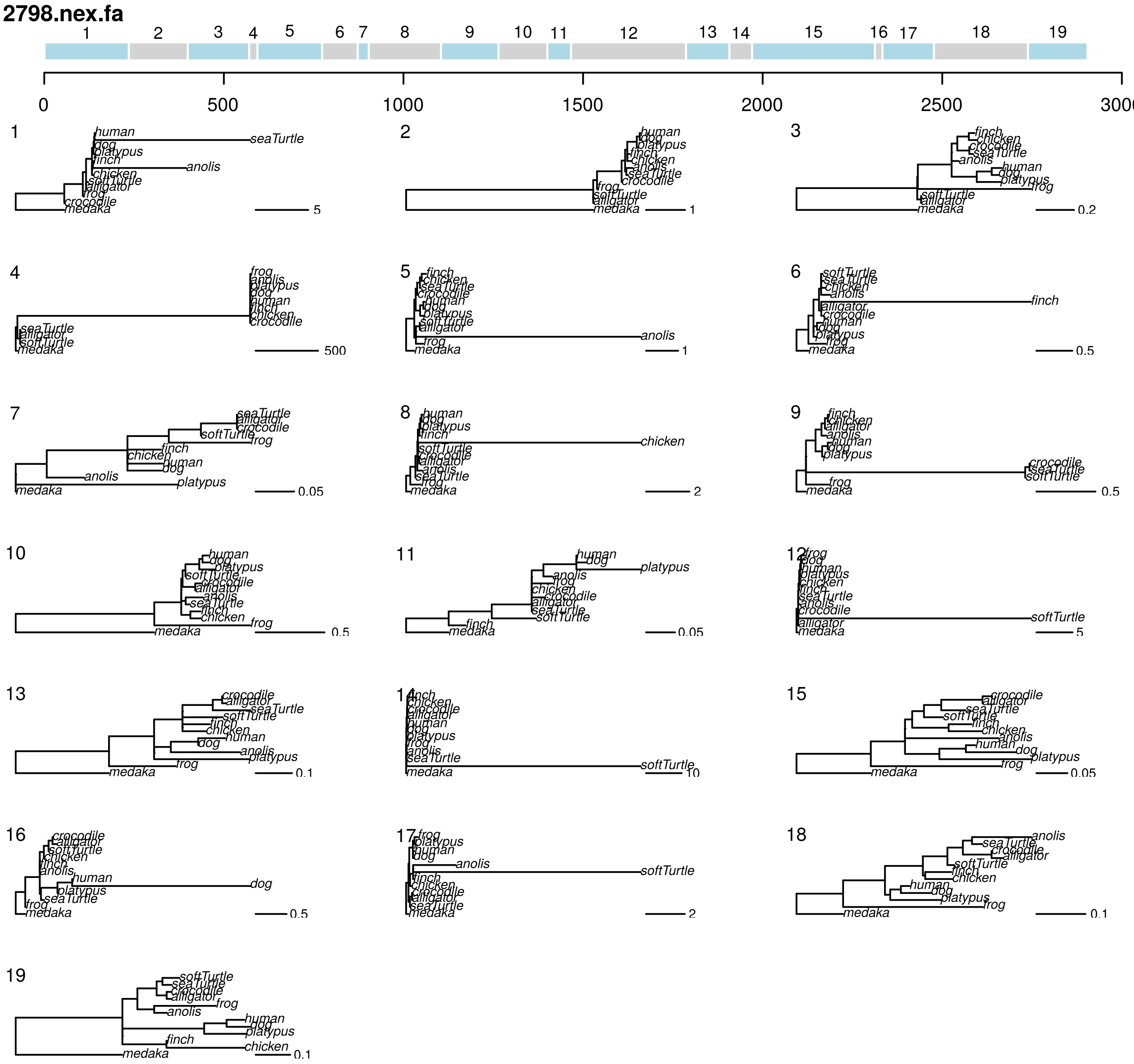


**Figure S11:** replicate one GARD analysis of 2798 under GTR+G. Tree topologies correspond to numbered segments, colors are arbitrary. Branch lengths and scale bar correspond to expected nucleotide substitutions per site.

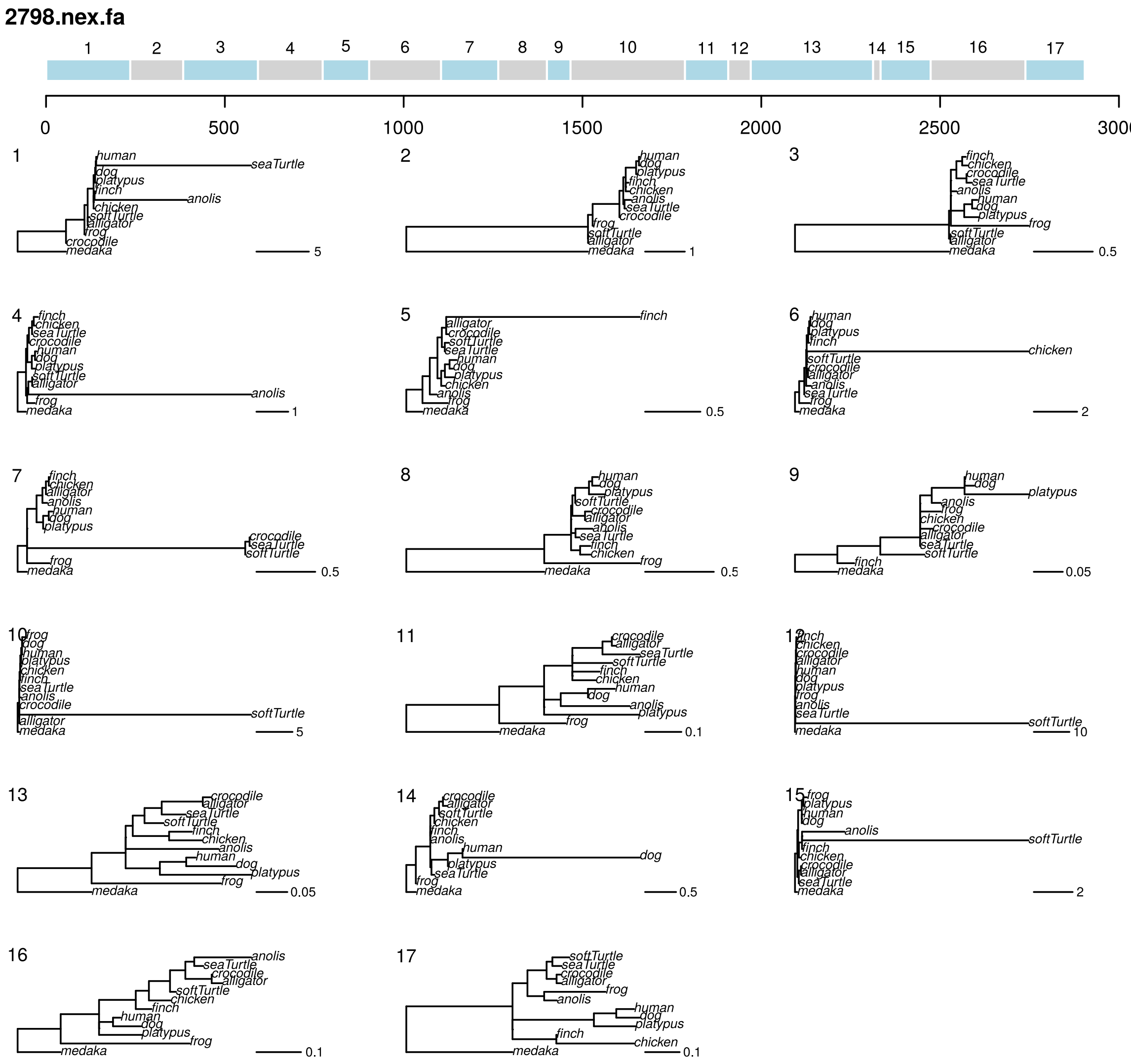


**Figure S12:** replicate two GARD analysis of 2798 under GTR+G. Tree topologies correspond to numbered segments, colors are arbitrary. Branch lengths and scale bar correspond to expected nucleotide substitutions per site.

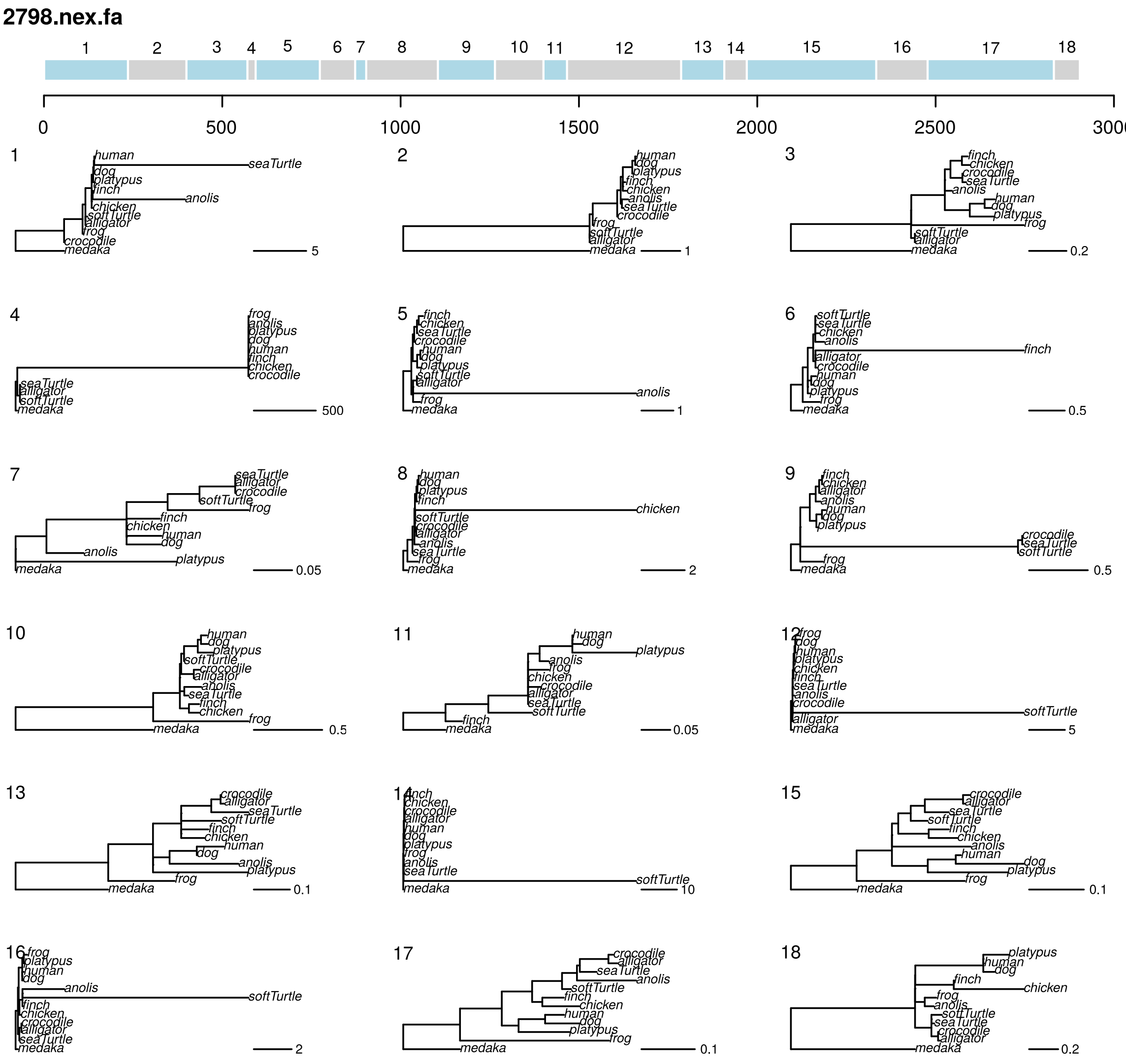


**Figure S13:** replicate three GARD analysis of 2798 under GTR+G. Tree topologies correspond to numbered segments, colors are arbitrary. Branch lengths and scale bar correspond to expected nucleotide substitutions per site.

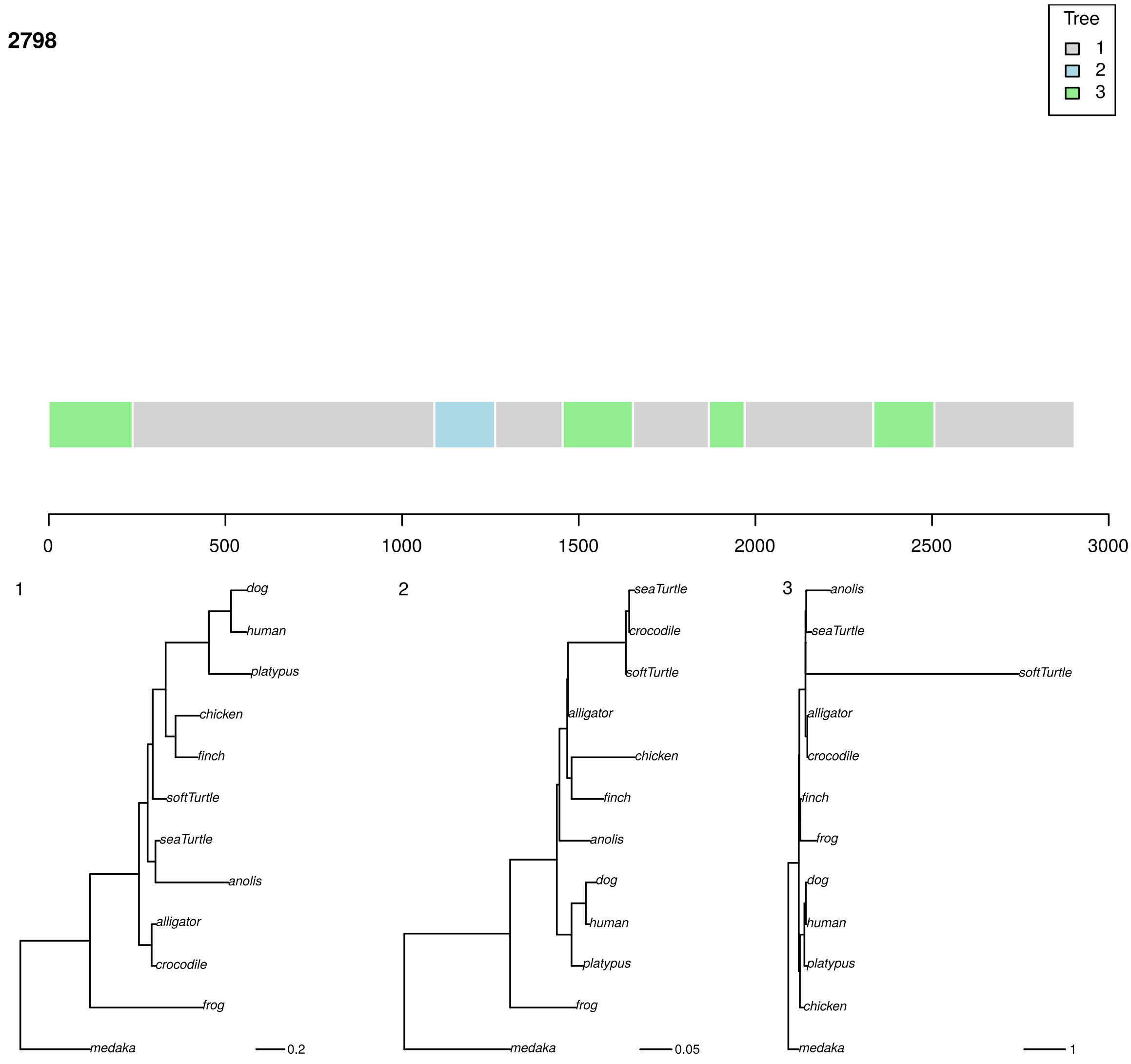


**Figure S14:** phyML_multi analysis of 2798 under TN93+G with three trees. Colors correspond to the best-supported tree for each segment. Branch lengths and scale bar correspond to expected nucleotide substitutions per site.

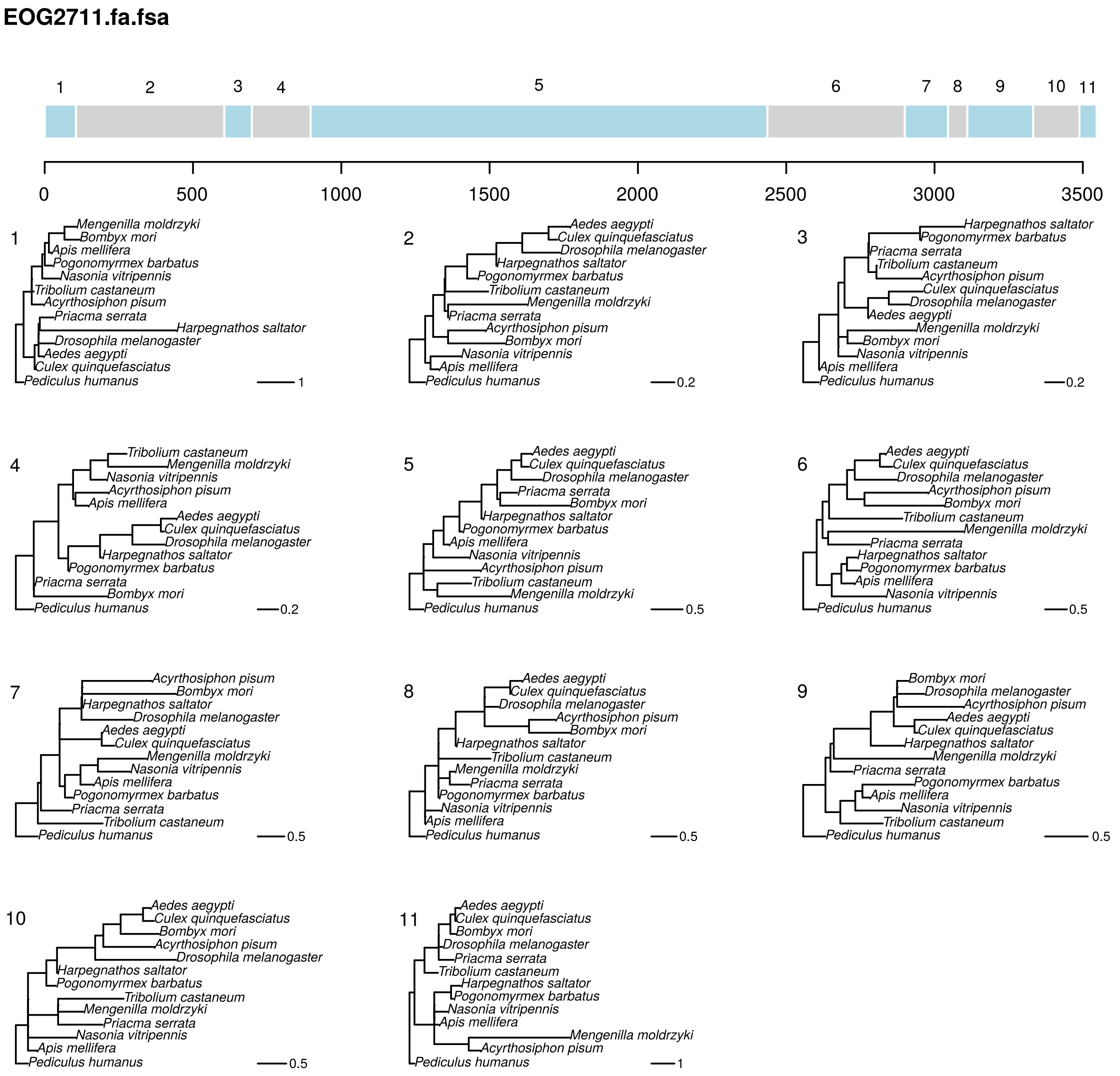


**Figure S15:** replicate one GARD analysis of an FSA realignment of EOG2711 under GTR+G. Tree topologies correspond to numbered segments, colors are arbitrary. Branch lengths and scale correspond to expected nucleotide substitutions per site.

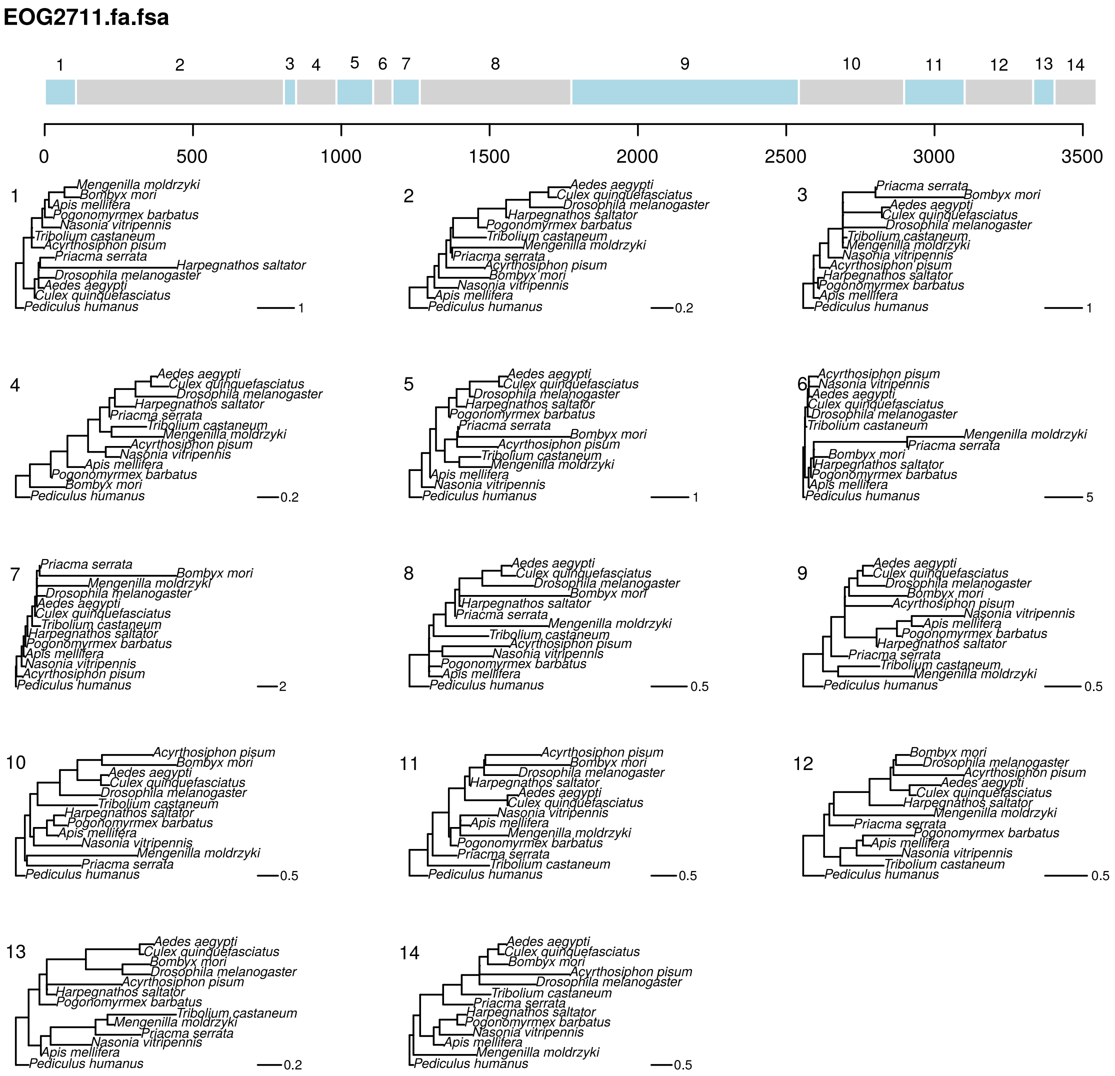


**Figure S16:** replicate two GARD analysis of an FSA realignment of EOG2711 under GTR+G. Tree topologies correspond to numbered segments, colors are arbitrary. Branch lengths and scale correspond to expected nucleotide substitutions per site.

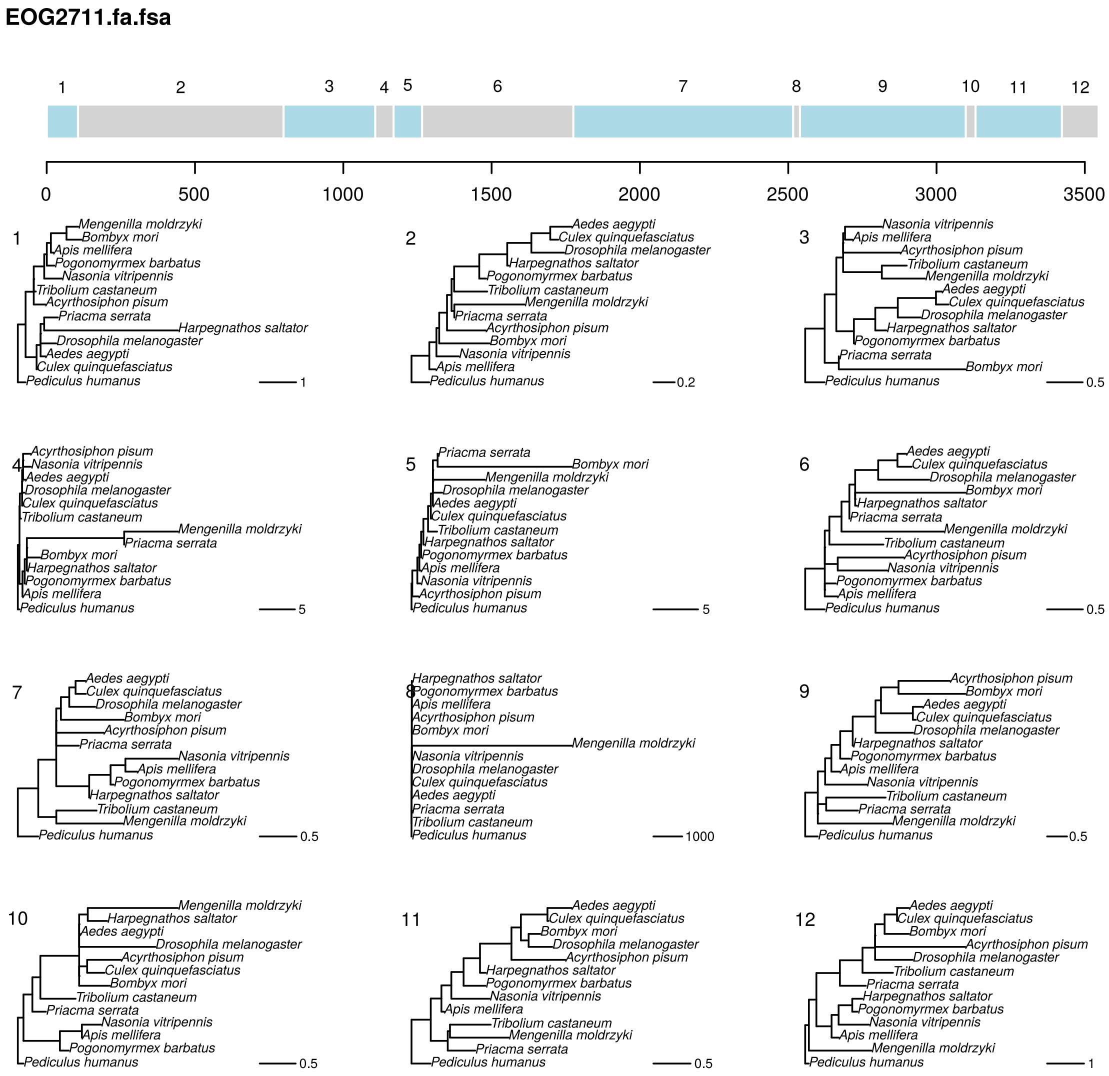


**Figure S17:** replicate three GARD analysis of an FSA realignment of EOG2711 under GTR+G. Tree topologies correspond to numbered segments, colors are arbitrary. Branch lengths and scale correspond to expected nucleotide substitutions per site.

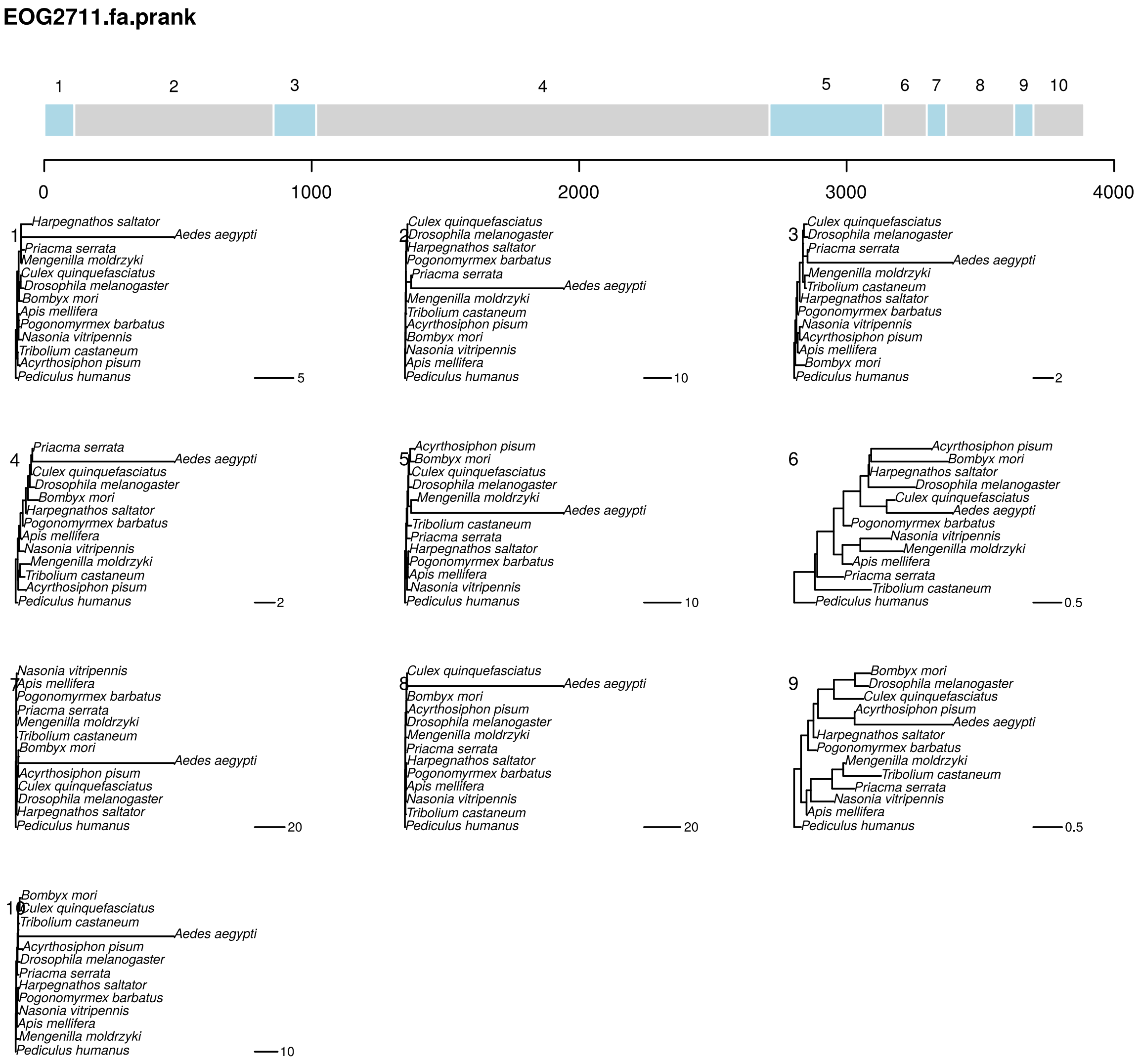


**Figure S18:** replicate one GARD analysis of a PRANK realignment of EOG2711 under GTR+G. Tree topologies correspond to numbered segments, colors are arbitrary. Branch lengths and scale correspond to expected nucleotide substitutions per site.

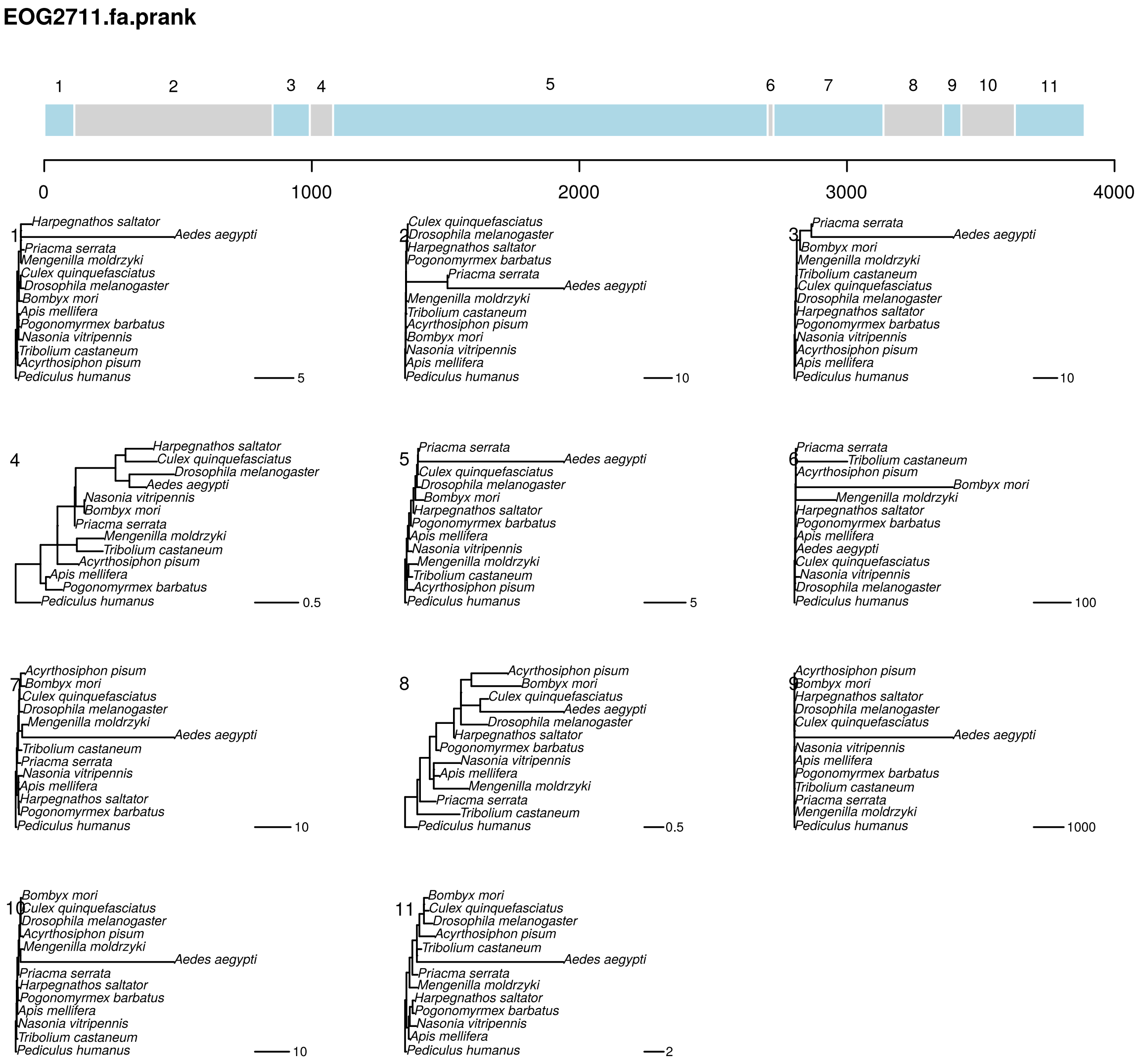


**Figure S19:** replicate two GARD analysis of a PRANK realignment of EOG2711 under GTR+G. Tree topologies correspond to numbered segments, colors are arbitrary. Branch lengths and scale correspond to expected nucleotide substitutions per site.

**EOG2711.fa.prank**

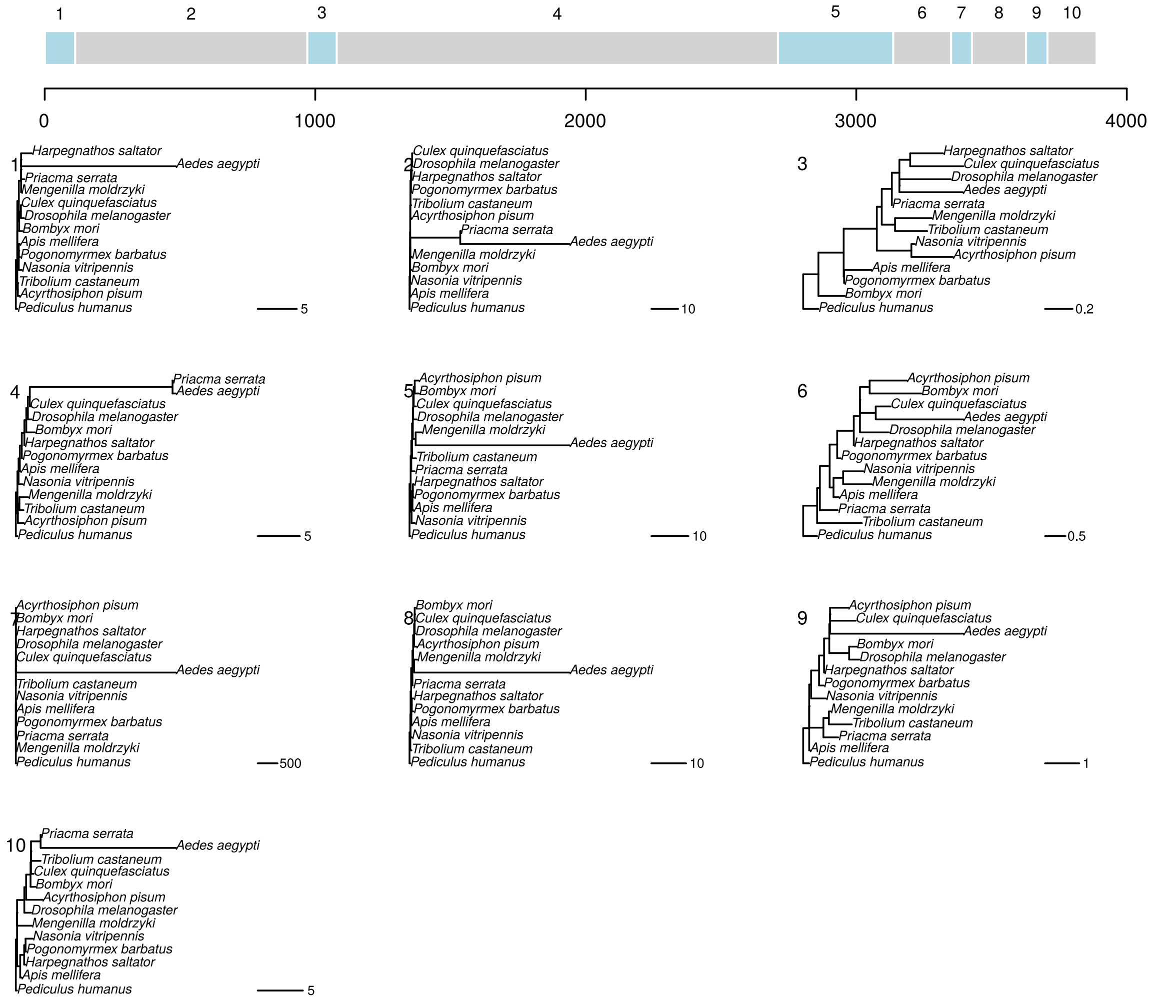


**Figure S20:** replicate three GARD analysis of a PRANK realignment of EOG2711 under GTR+G. Tree topologies correspond to numbered segments, colors are arbitrary. Branch lengths and scale correspond to expected nucleotide substitutions per site.

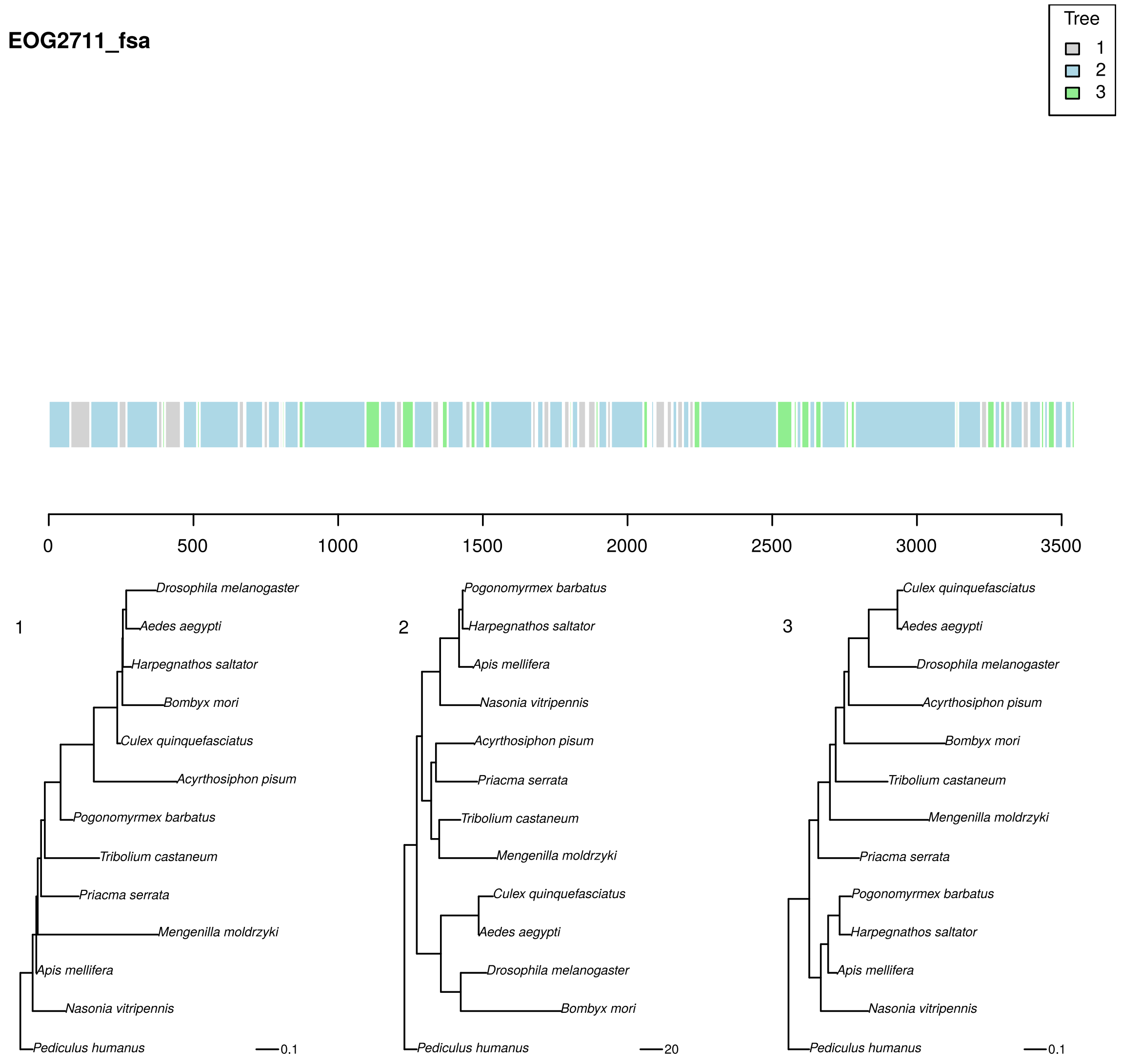


**Figure S21:** phyML_multi analysis of an FSA realignment of EOG2711 under TN93+G with three trees. Colors correspond to the best-supported tree for each segment. Branch lengths and scale bar correspond to expected nucleotide substitutions per site.

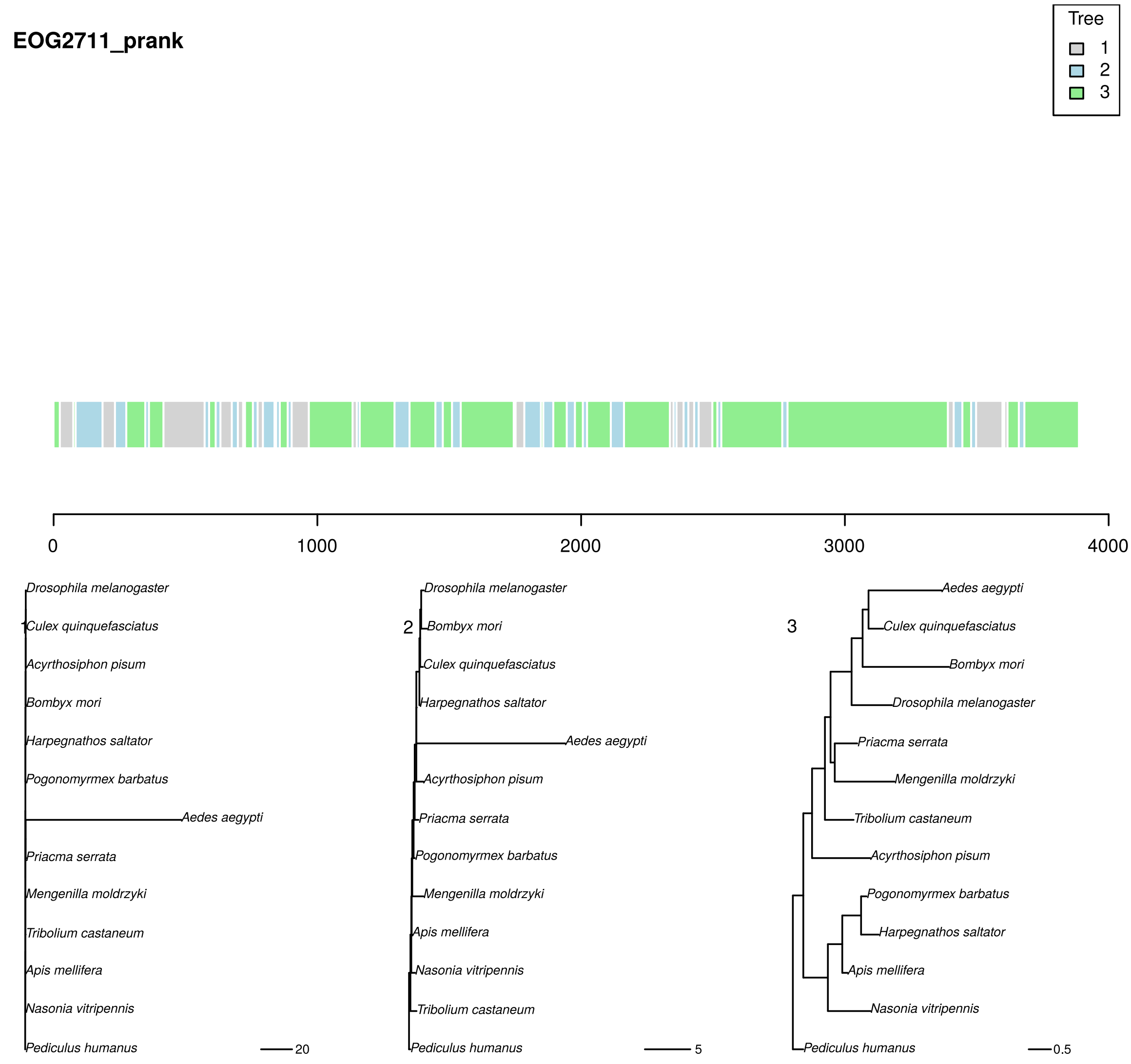


**Figure S22:** phyML_multi analysis of a PRANK realignment of EOG2711 under TN93+G with three trees. Colors correspond to the best-supported tree for each segment. Branch lengths and scale bar correspond to expected nucleotide substitutions per site.

**ENSG00000074803_SLC12A1_rhinoceros_added_2N.fasta.fsa**

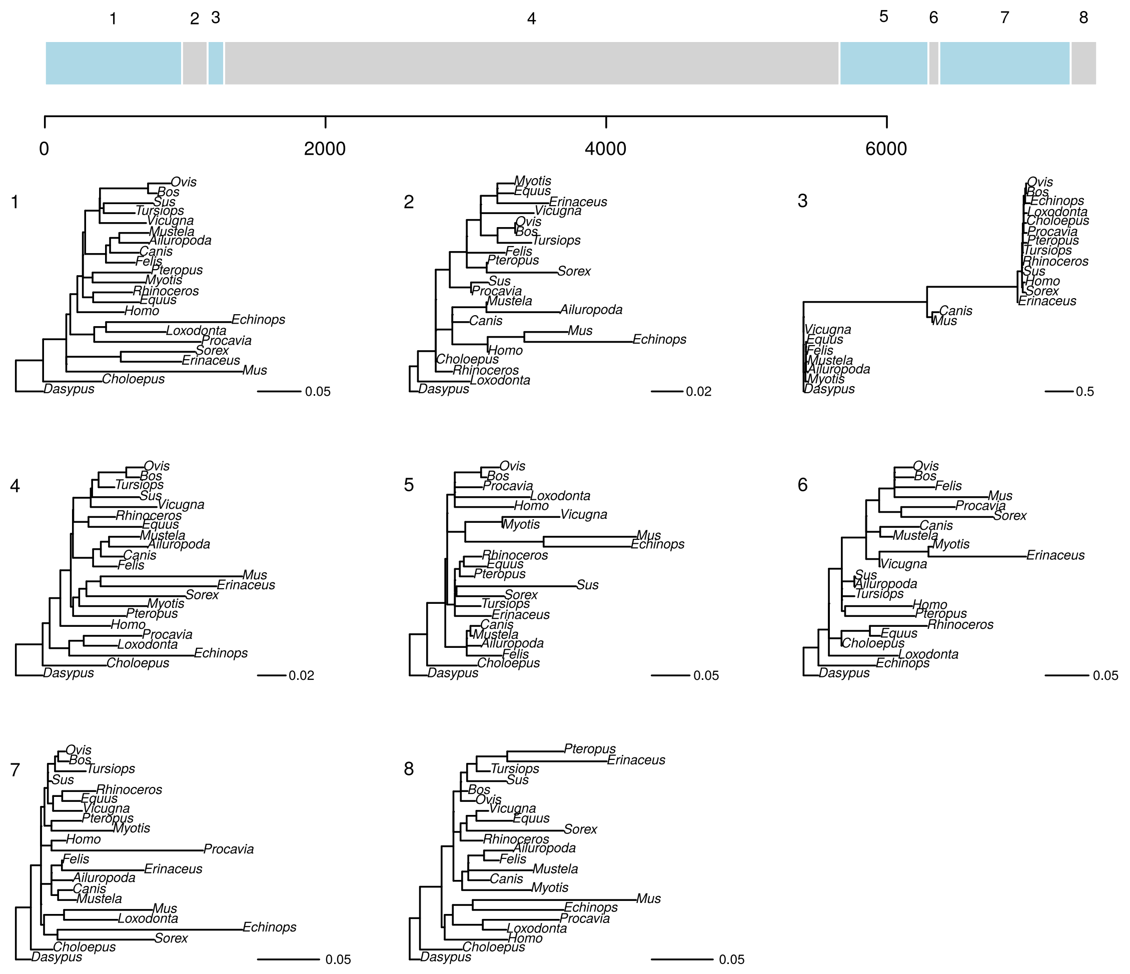


**Figure S23:** replicate one GARD analysis of an FSA realignment of ENSG00000074803 under GTR+G. Tree topologies correspond to numbered segments, colors are arbitrary. Branch lengths and scale correspond to expected nucleotide substitutions per site.

**ENSG00000074803_SLC12A1_rhinoceros_added_2N.fasta.fsa**

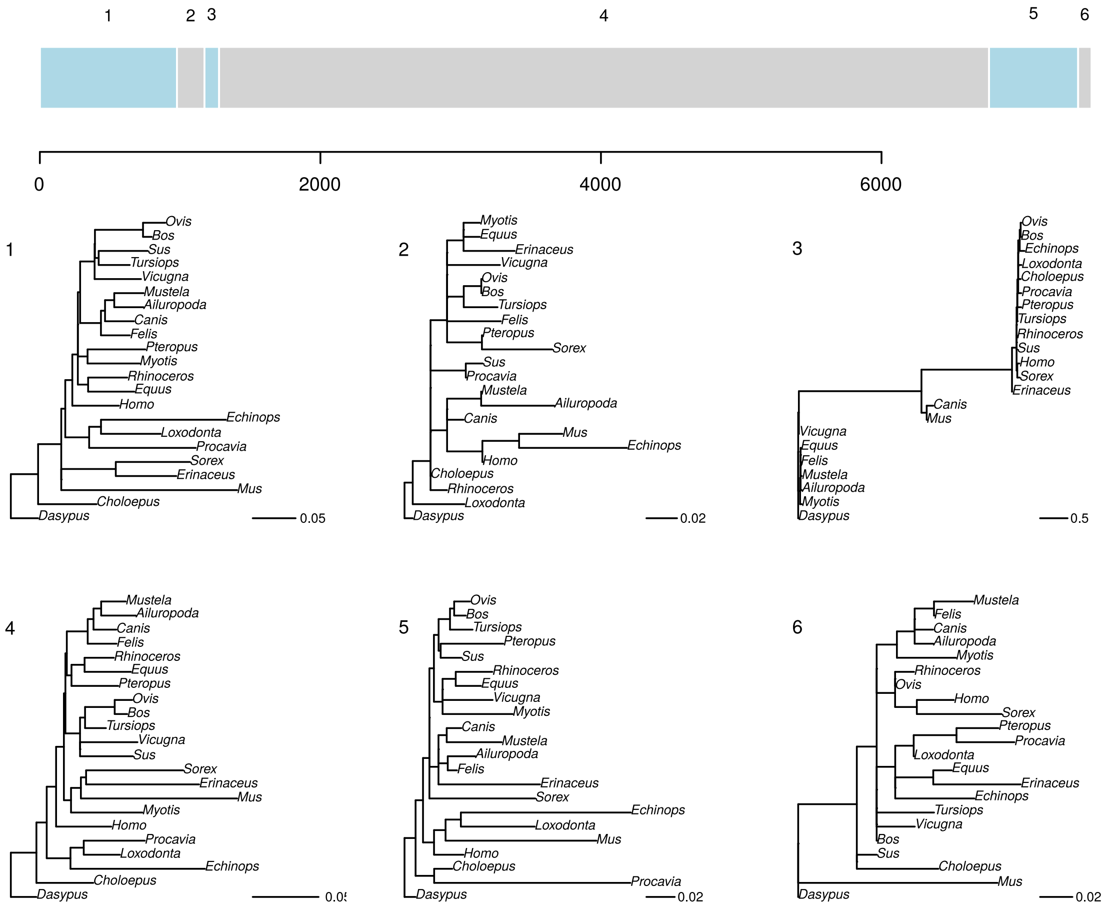


**Figure S24:** replicate two GARD analysis of an FSA realignment of ENSG00000074803 under GTR+G. Tree topologies correspond to numbered segments, colors are arbitrary. Branch lengths and scale correspond to expected nucleotide substitutions per site.

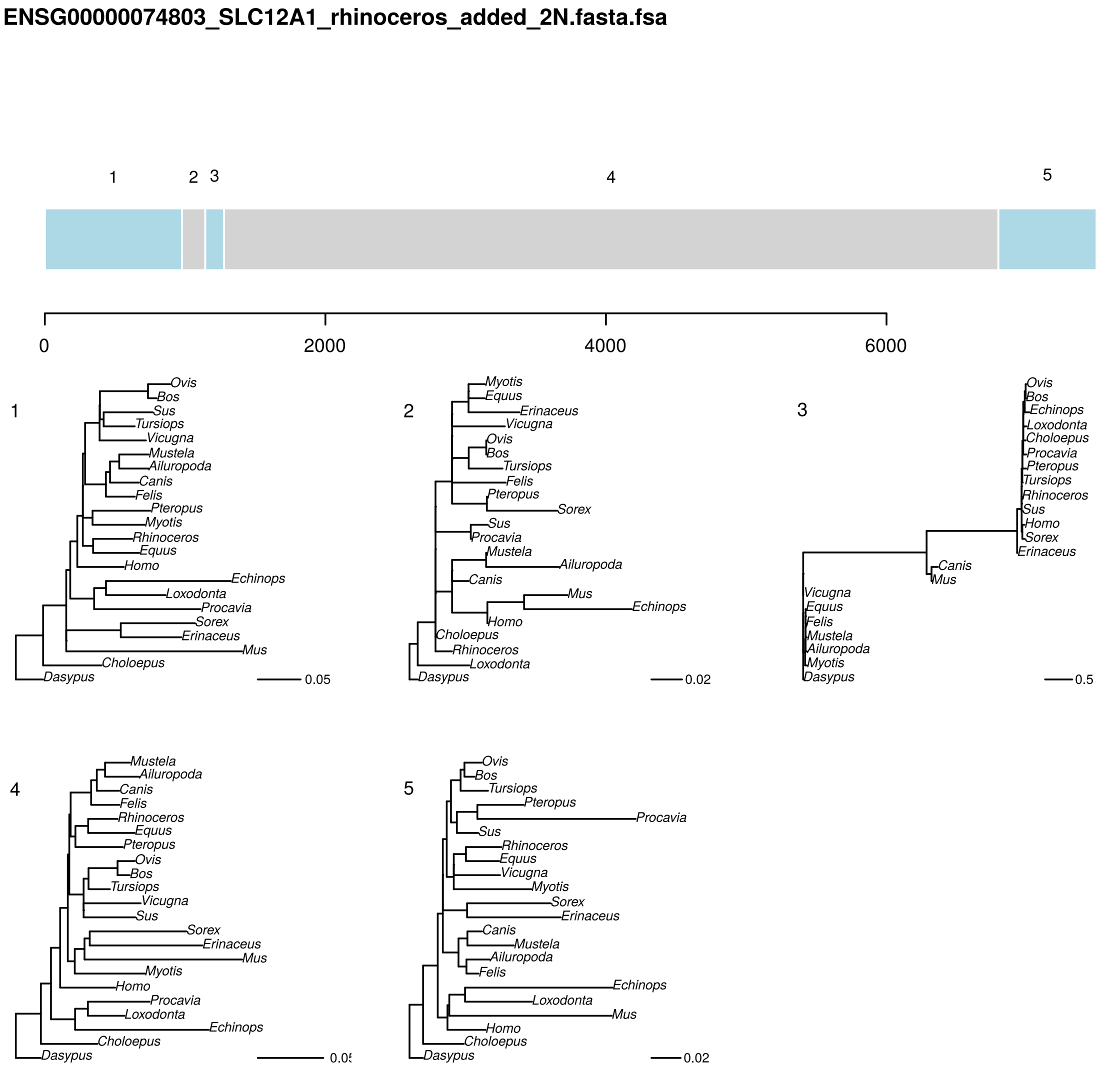


**Figure S25:** replicate three GARD analysis of an FSA realignment of ENSG00000074803 under GTR+G. Tree topologies correspond to numbered segments, colors are arbitrary. Branch lengths and scale correspond to expected nucleotide substitutions per site.

**ENSG00000074803_SLC12A1_rhinoceros_added_2N.fasta.prank**

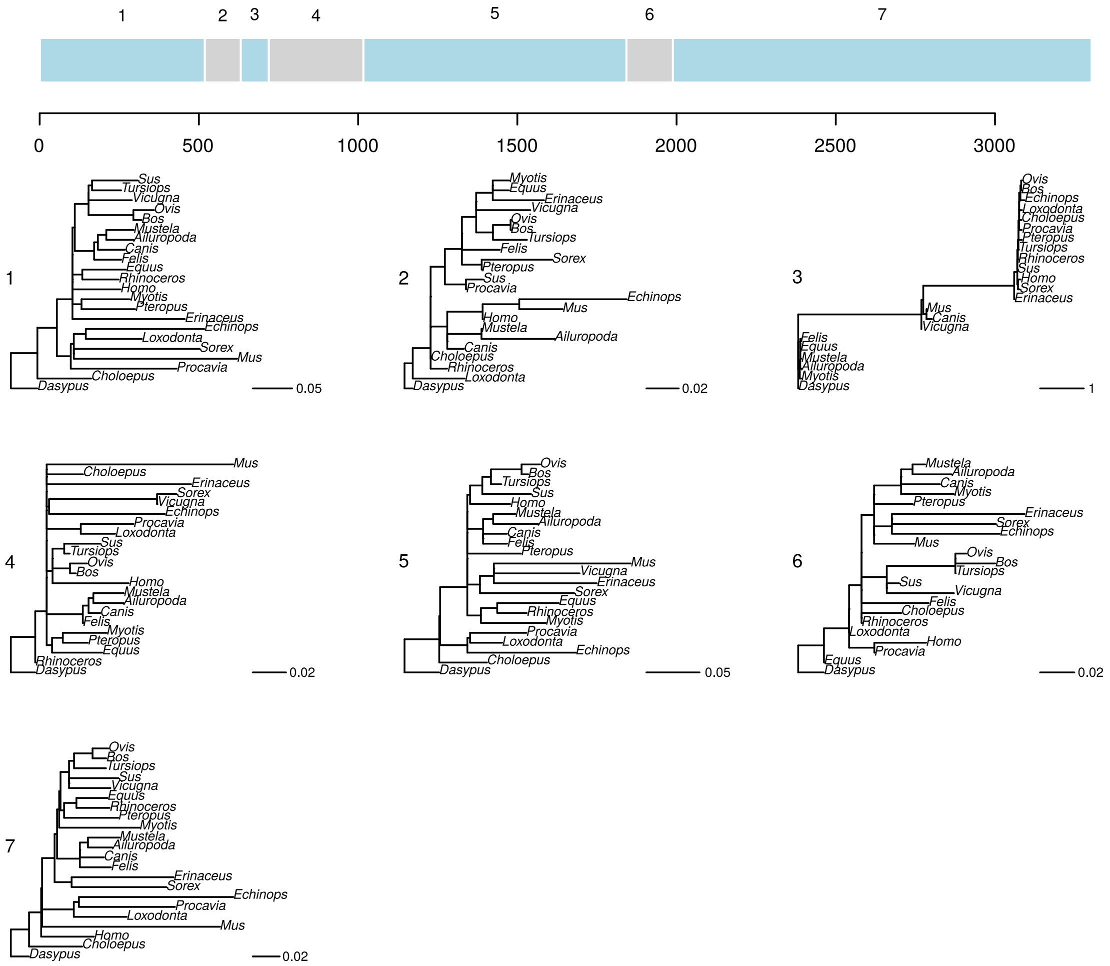


**Figure S26:** replicate one GARD analysis of a PRANK realignment of ENSG00000074803 under GTR+G. Tree topologies correspond to numbered segments, colors are arbitrary. Branch lengths and scale correspond to expected nucleotide substitutions per site.

### ENSG00000074803_SLC12A1_rhinoceros_added_2N.fasta.prank

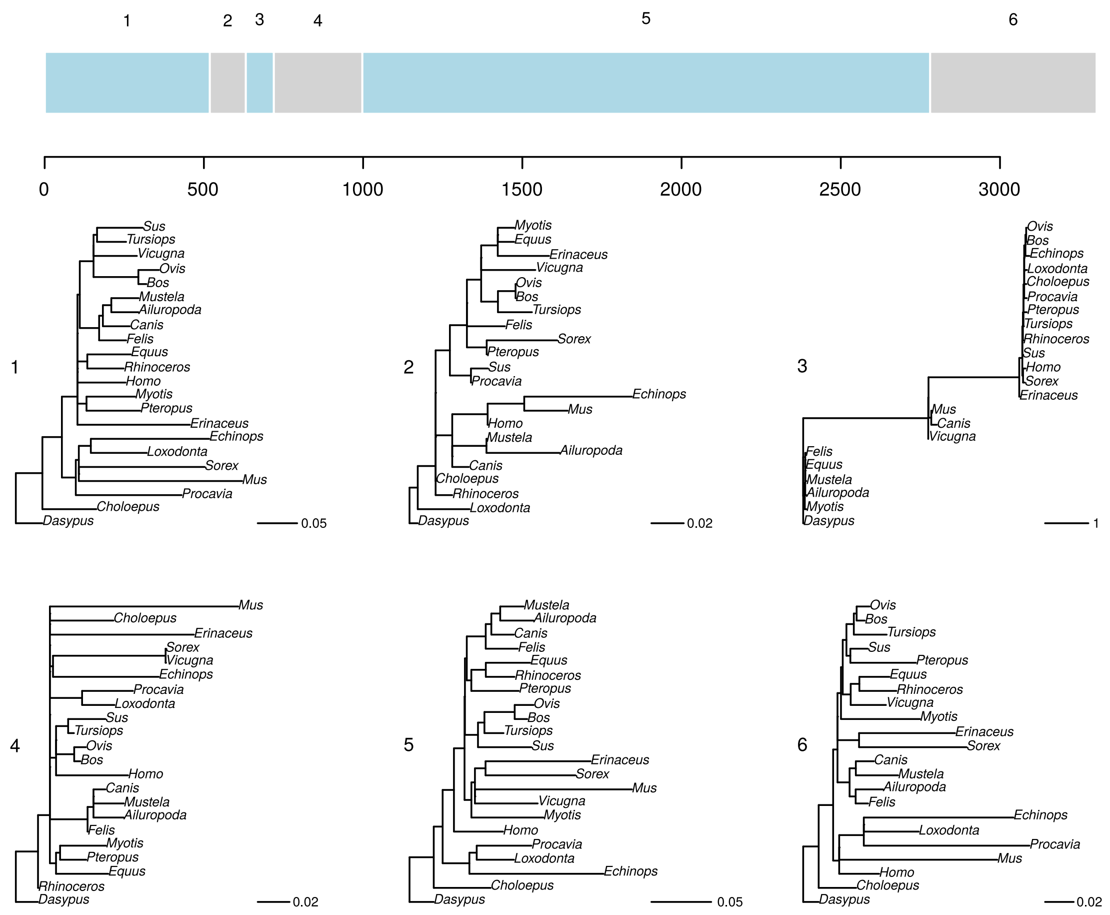


**Figure S27:** replicate two GARD analysis of a PRANK realignment of ENSG00000074803 under GTR+G. Tree topologies correspond to numbered segments, colors are arbitrary. Branch lengths and scale correspond to expected nucleotide substitutions per site.

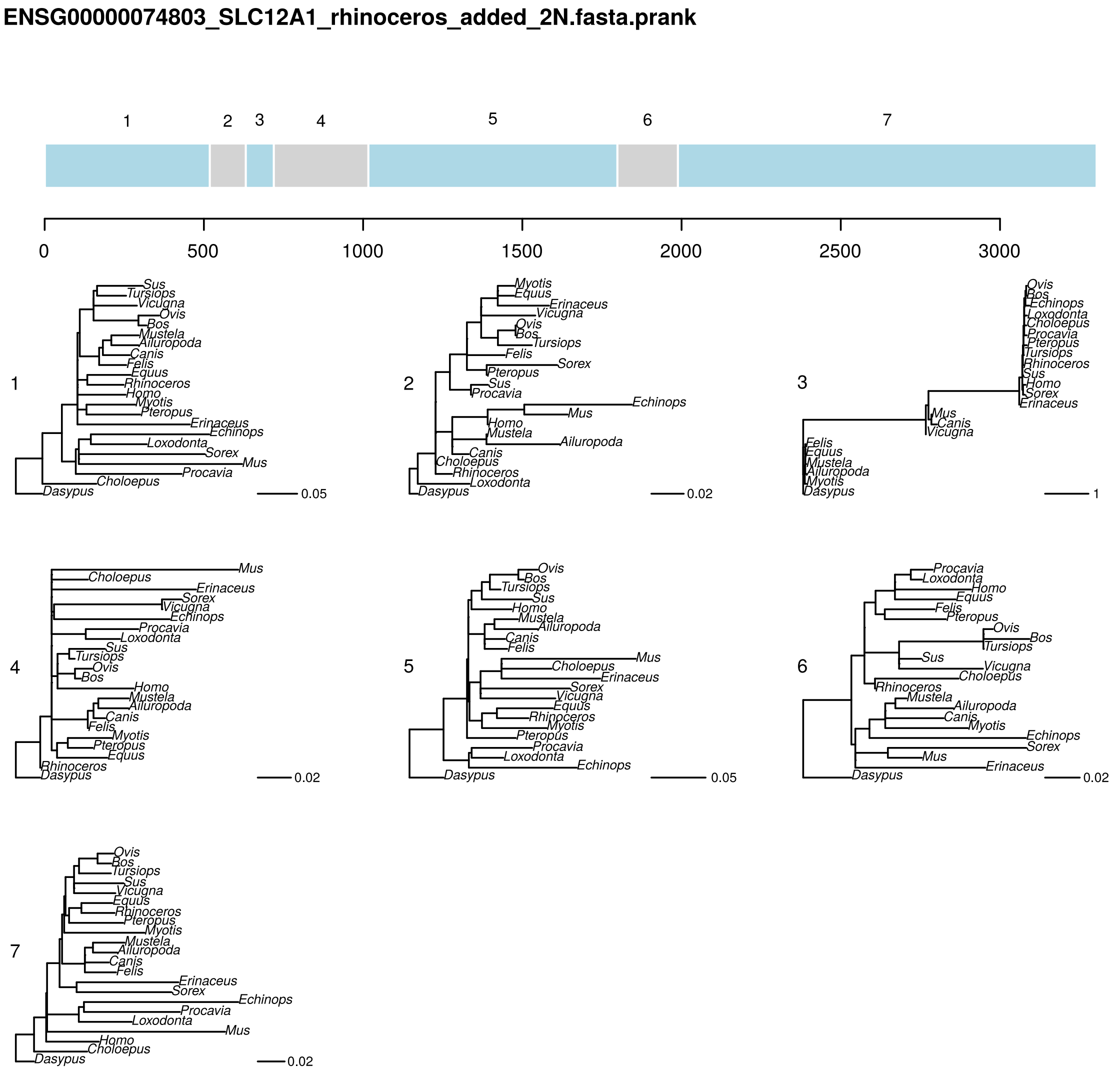


**Figure S28:** replicate three GARD analysis of a PRANK realignment of ENSG00000074803 under GTR+G. Tree topologies correspond to numbered segments, colors are arbitrary. Branch lengths and scale correspond to expected nucleotide substitutions per site.

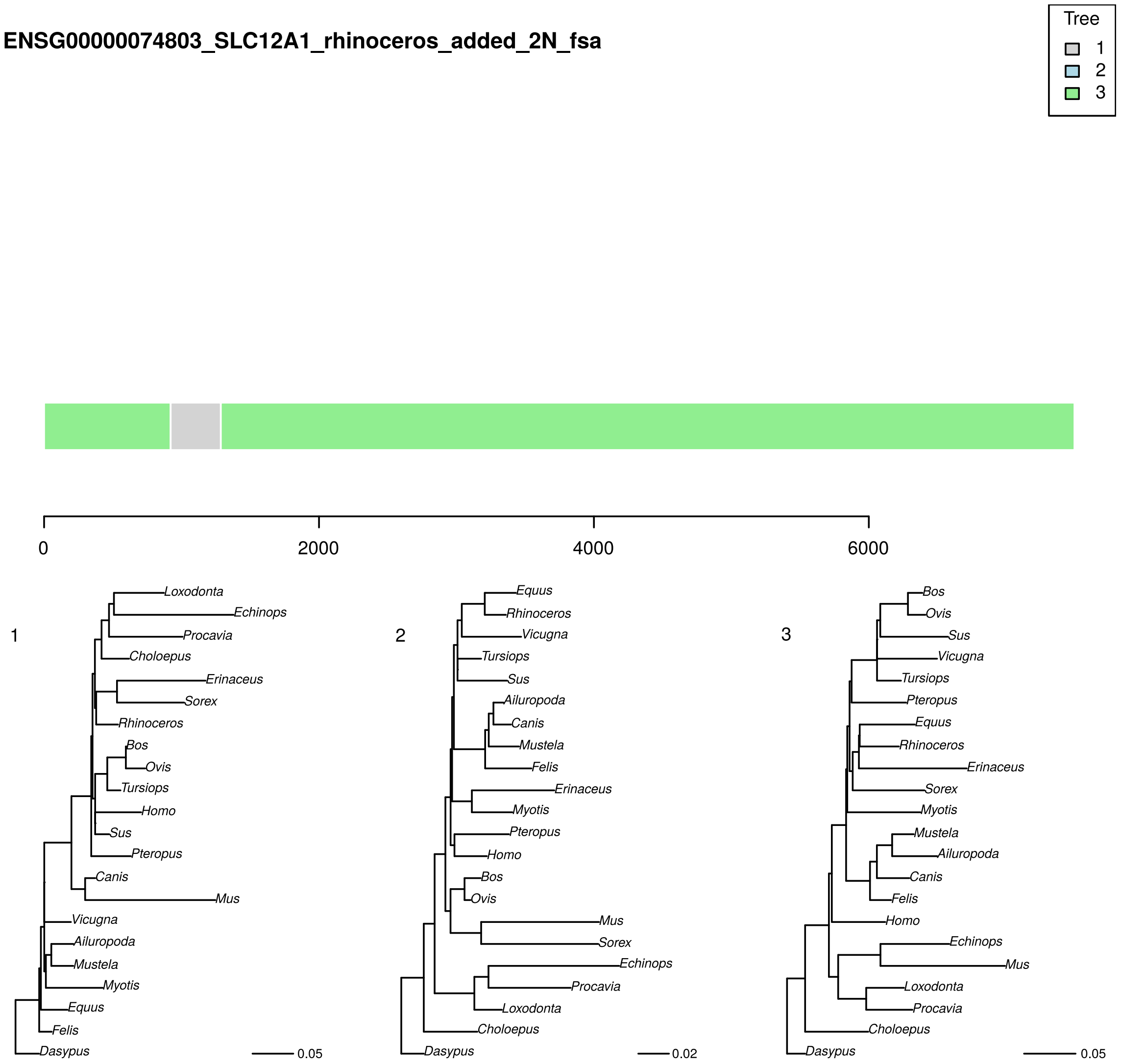


**Figure S29:** phyML_multi analysis of an FSA realignment of ENSG00000074803 under TN93+G with three trees. Colors correspond to the best-supported tree for each segment. Branch lengths and scale bar correspond to expected nucleotide substitutions per site.

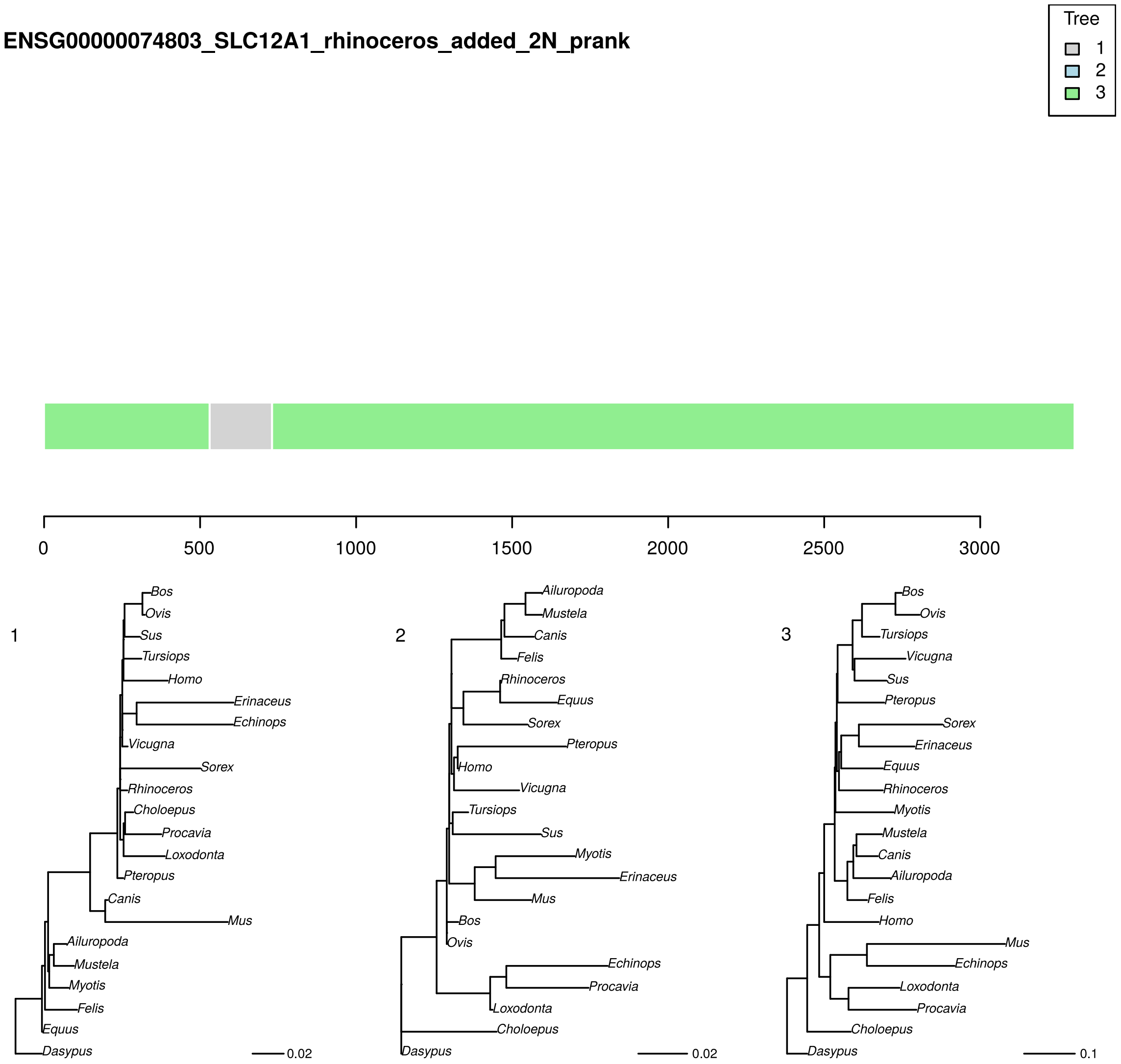


**Figure S30:** phyML_multi analysis of a PRANK realignment of ENSG00000074803 under TN93+G with three trees. Colors correspond to the best-supported tree for each segment. Branch lengths and scale bar correspond to expected nucleotide substitutions per site.

**131.fasta.fsa**

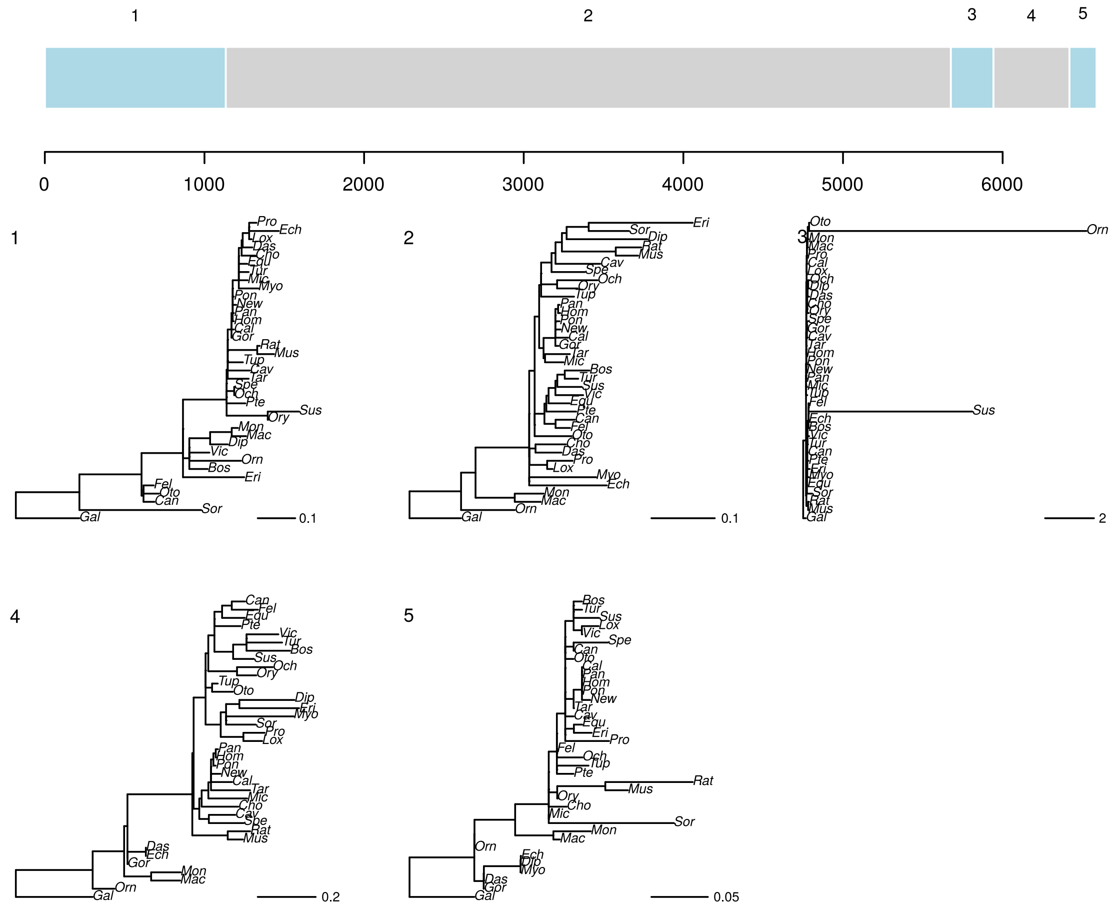


**Figure S31:** replicate one GARD analysis of an FSA realignment of 131 under GTR+G. Tree topologies correspond to numbered segments, colors are arbitrary. Branch lengths and scale correspond to expected nucleotide substitutions per site.

**131.fasta.fsa**

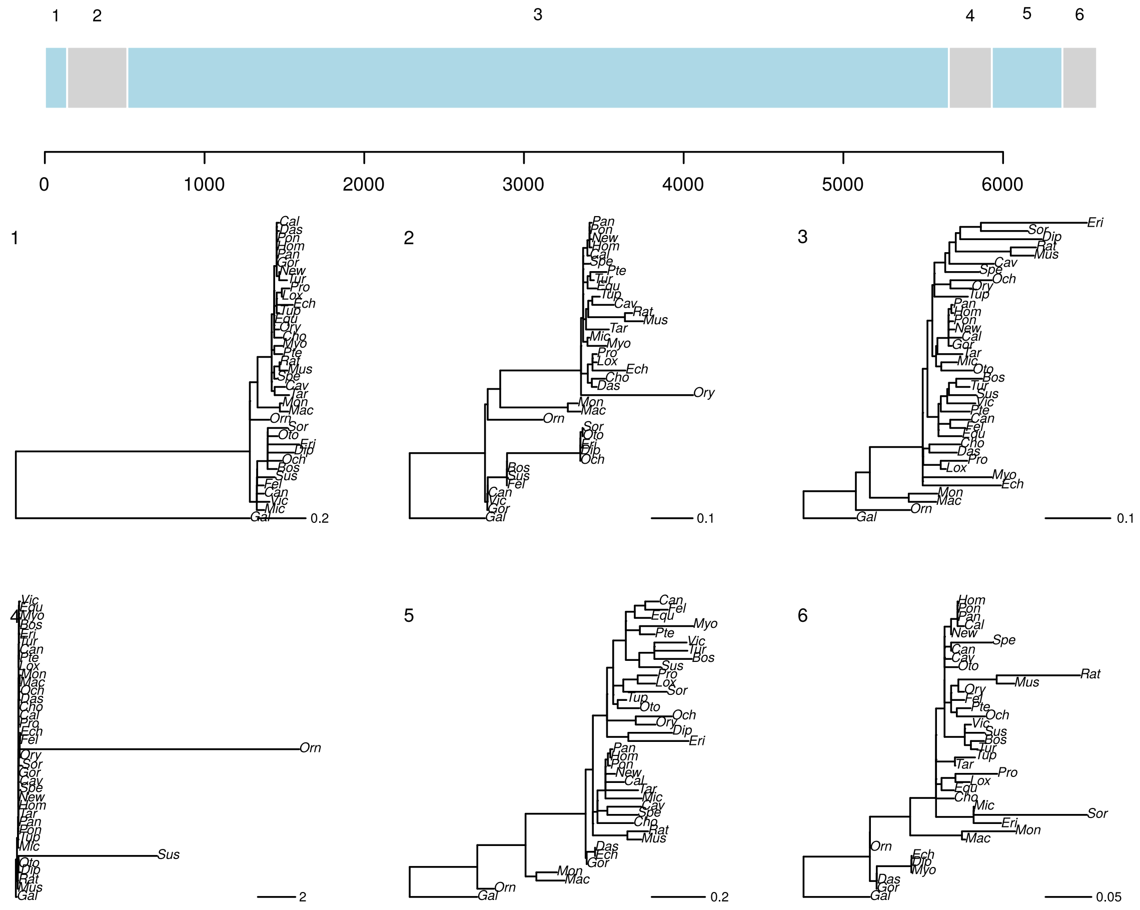


**Figure S32:** replicate two GARD analysis of an FSA realignment of 131 under GTR+G. Tree topologies correspond to numbered segments, colors are arbitrary. Branch lengths and scale correspond to expected nucleotide substitutions per site.

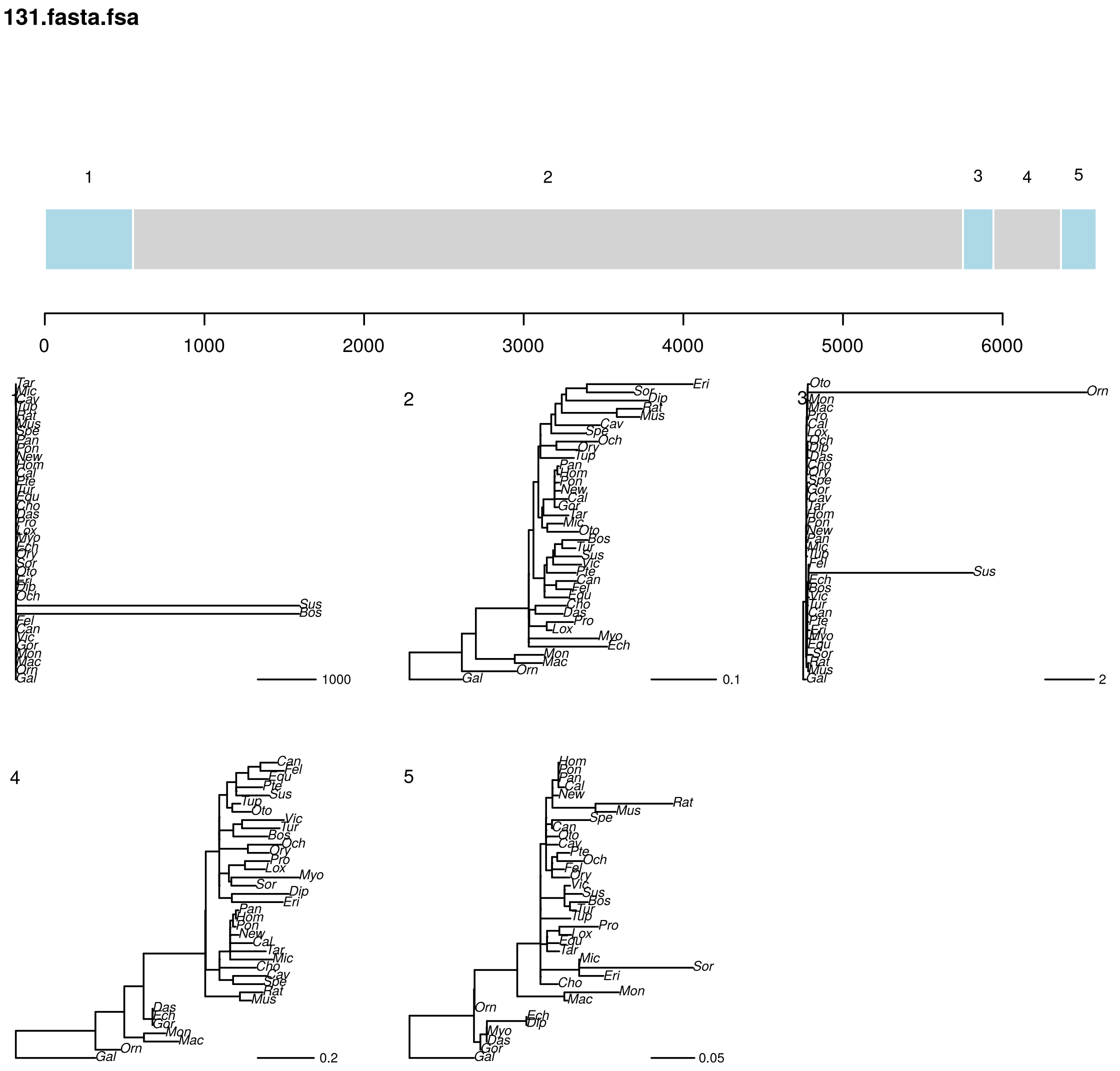


**Figure S33:** replicate three GARD analysis of an FSA realignment of 131 under GTR+G. Tree topologies correspond to numbered segments, colors are arbitrary. Branch lengths and scale correspond to expected nucleotide substitutions per site.

**131.fasta.prank**

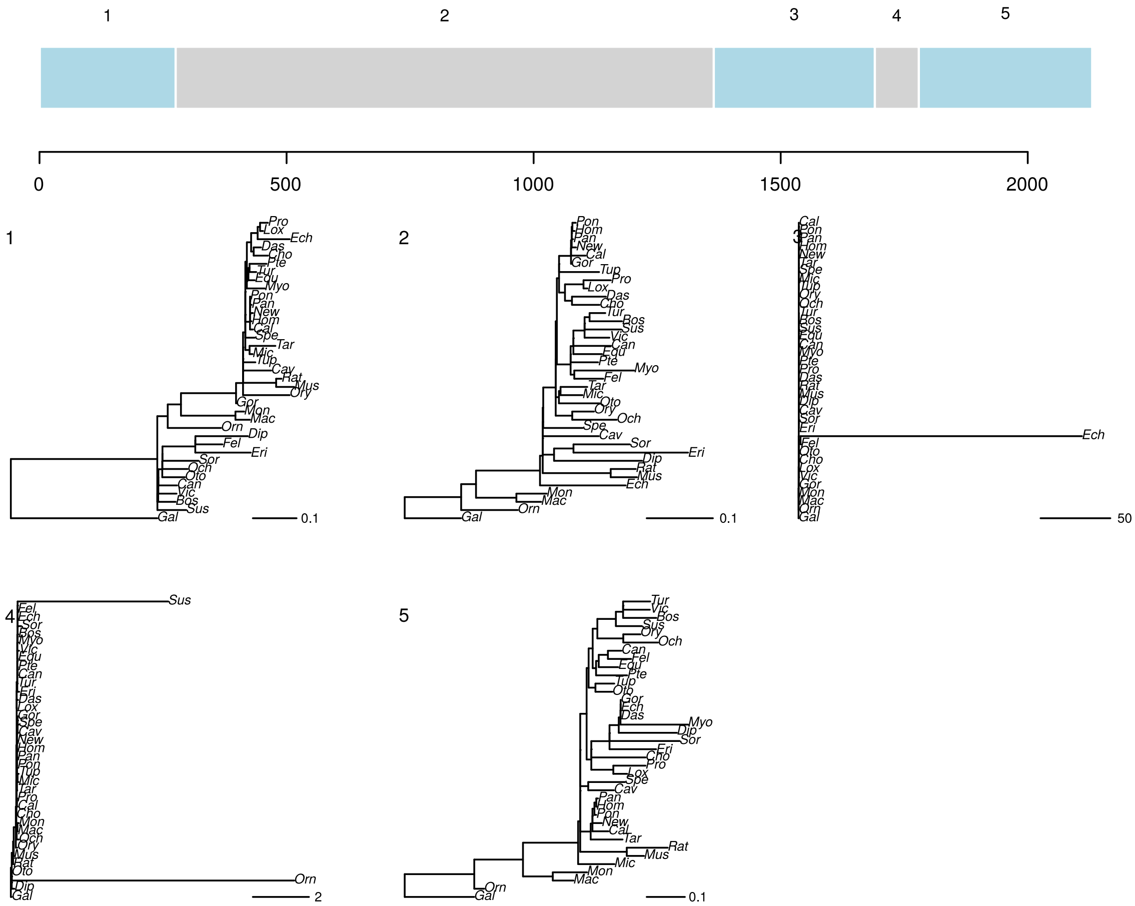


**Figure S34:** replicate one GARD analysis of a PRANK realignment of 131 under GTR+G. Tree topologies correspond to numbered segments, colors are arbitrary. Branch lengths and scale correspond to expected nucleotide substitutions per site.

**131.fasta.prank**

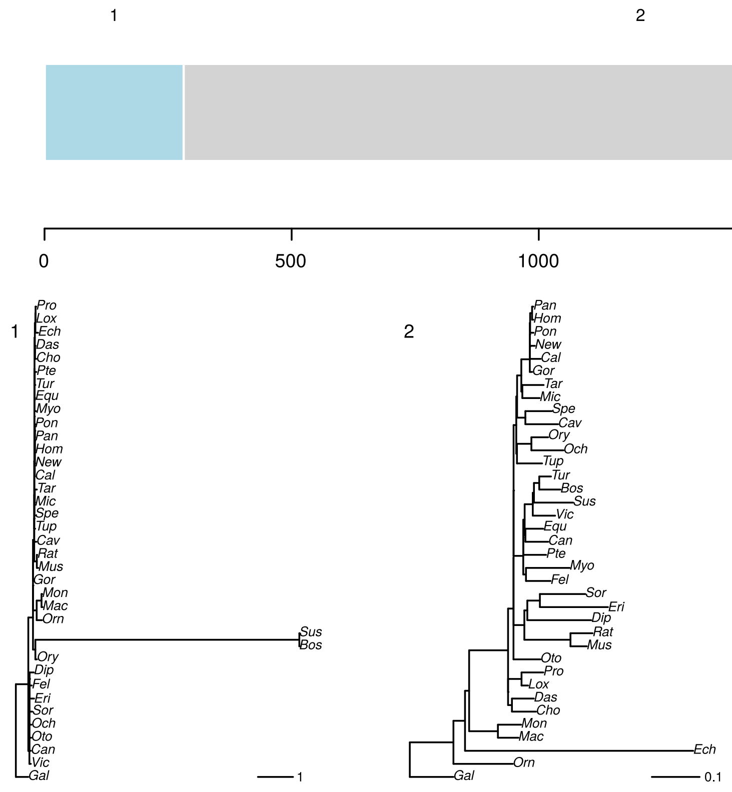


**Figure S35:** replicate two GARD analysis of a PRANK realignment of 131 under GTR+G. Tree topologies correspond to numbered segments, colors are arbitrary. Branch lengths and scale correspond to expected nucleotide substitutions per site.

**131.fasta.prank**

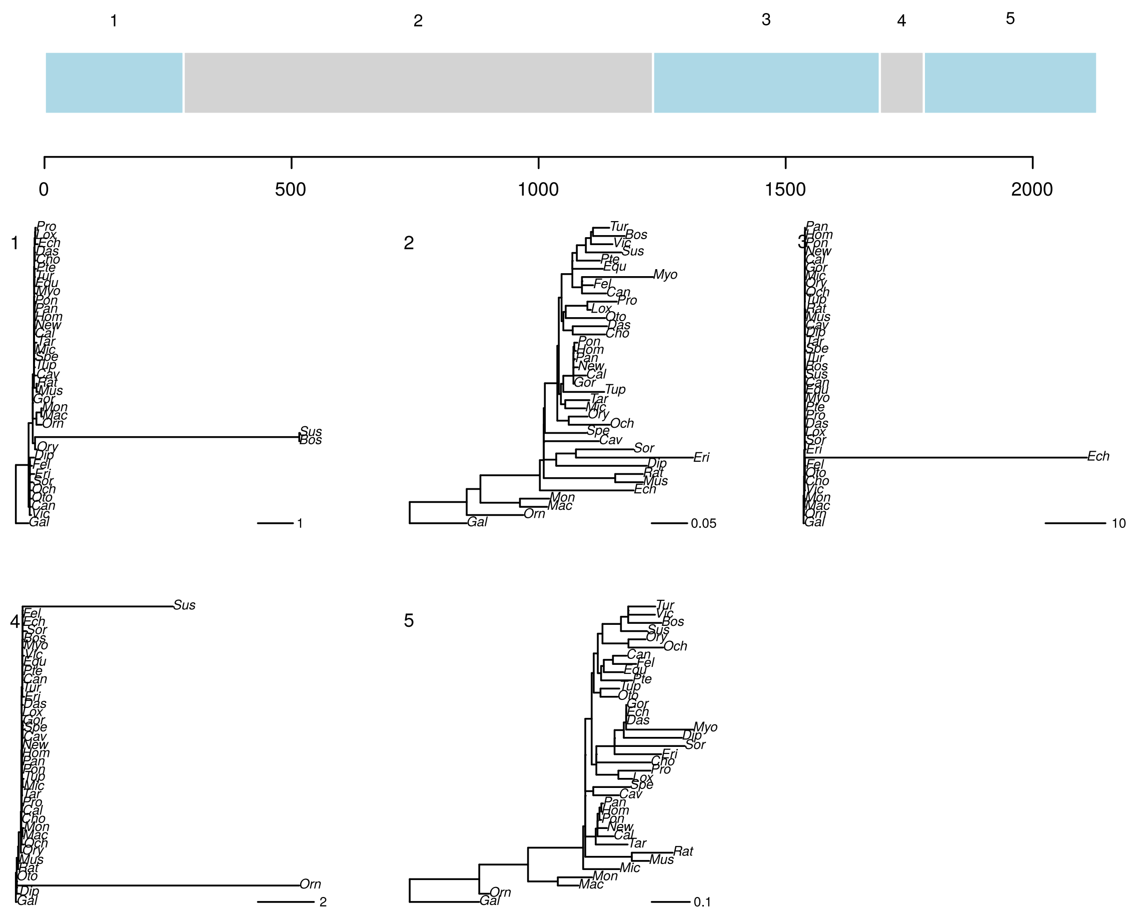


**Figure S36:** replicate three GARD analysis of a PRANK realignment of 131 under GTR+G. Tree topologies correspond to numbered segments, colors are arbitrary. Branch lengths and scale correspond to expected nucleotide substitutions per site.

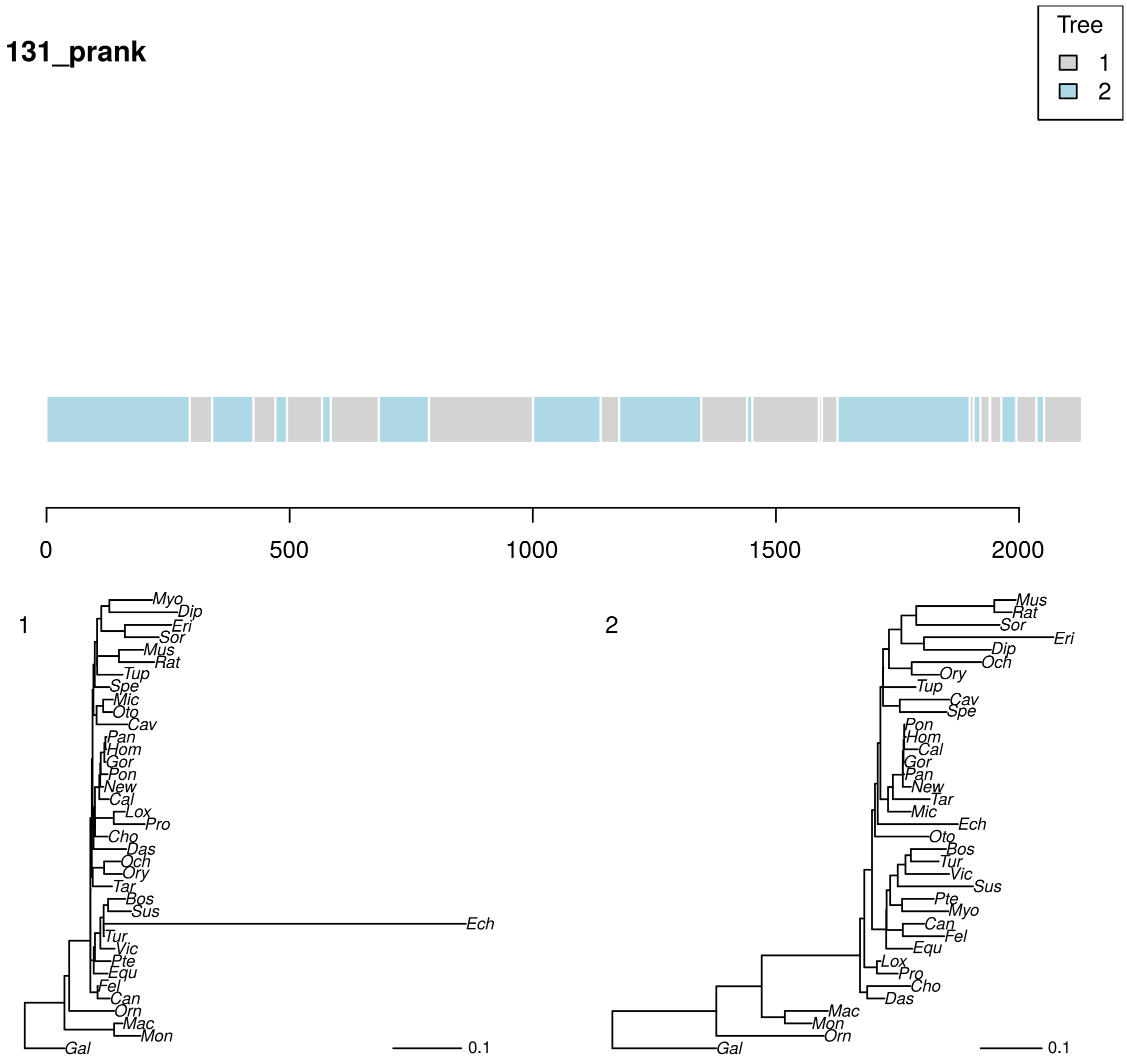


**Figure S37:** phyML_multi analysis of a PRANK realignment of 131 under TN93+G with two trees. Colors correspond to the best-supported tree for each segment. Branch lengths and scale bar correspond to expected nucleotide substitutions per site.

**2798.nex.fa**

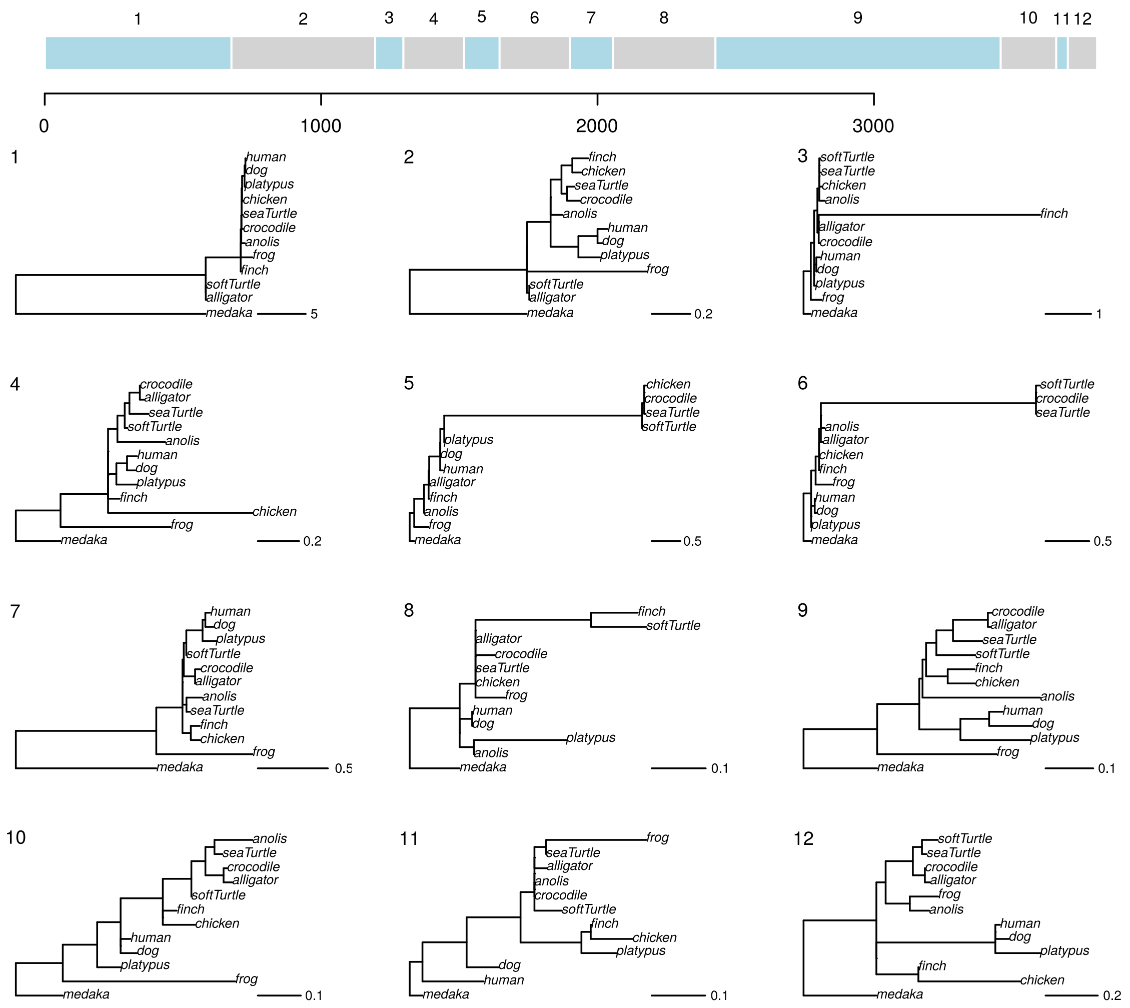


**Figure S38:** replicate one GARD analysis of an FSA realignment of 2798 under GTR+G. Tree topologies correspond to numbered segments, colors are arbitrary. Branch lengths and scale correspond to expected nucleotide substitutions per site.

**2798.nex.fa**

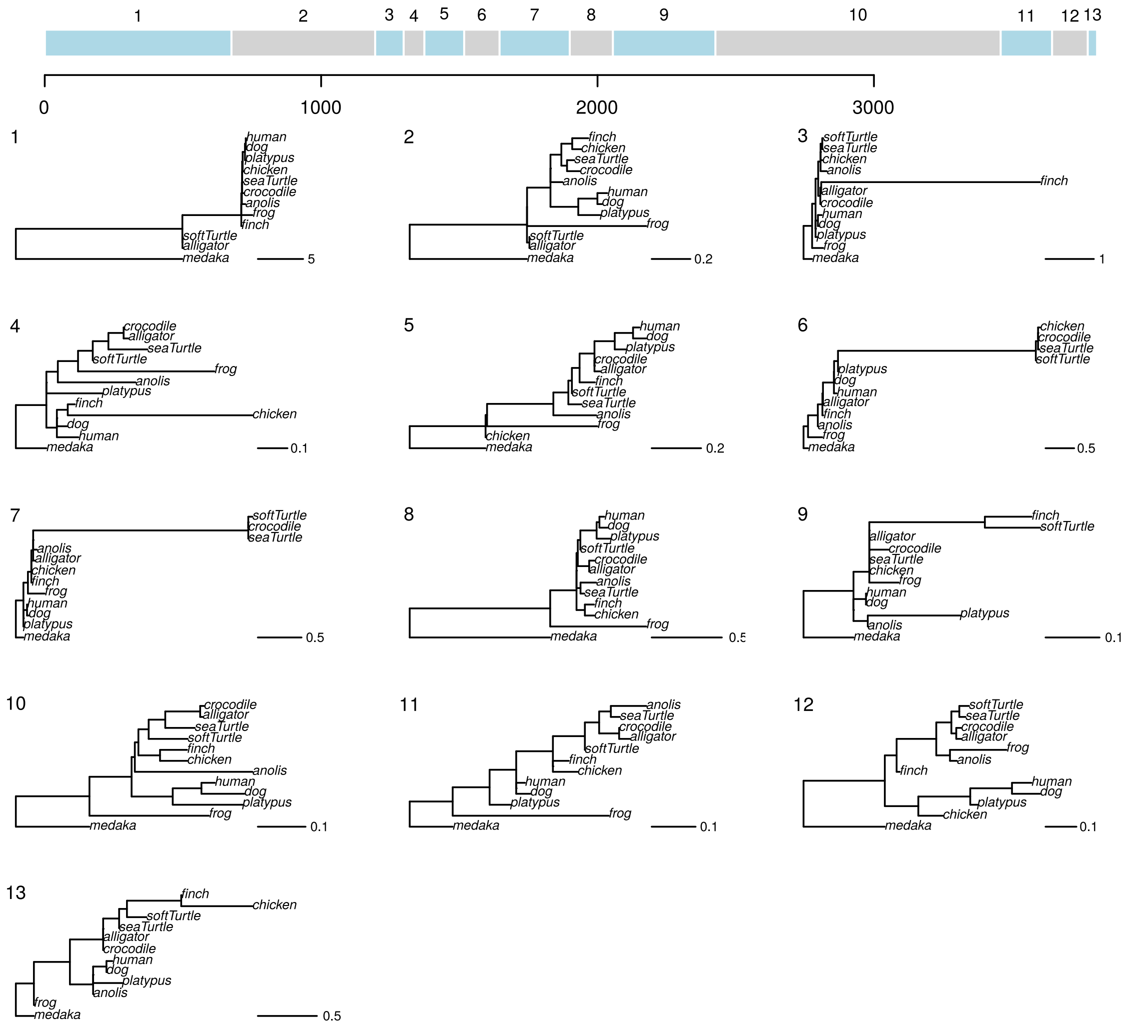

**Figure S39:** replicate two GARD analysis of an FSA realignment of 2798 under GTR+G. Tree topologies correspond to numbered segments, colors are arbitrary. Branch lengths and scale correspond to expected nucleotide substitutions per site.

**2798.nex.fa**

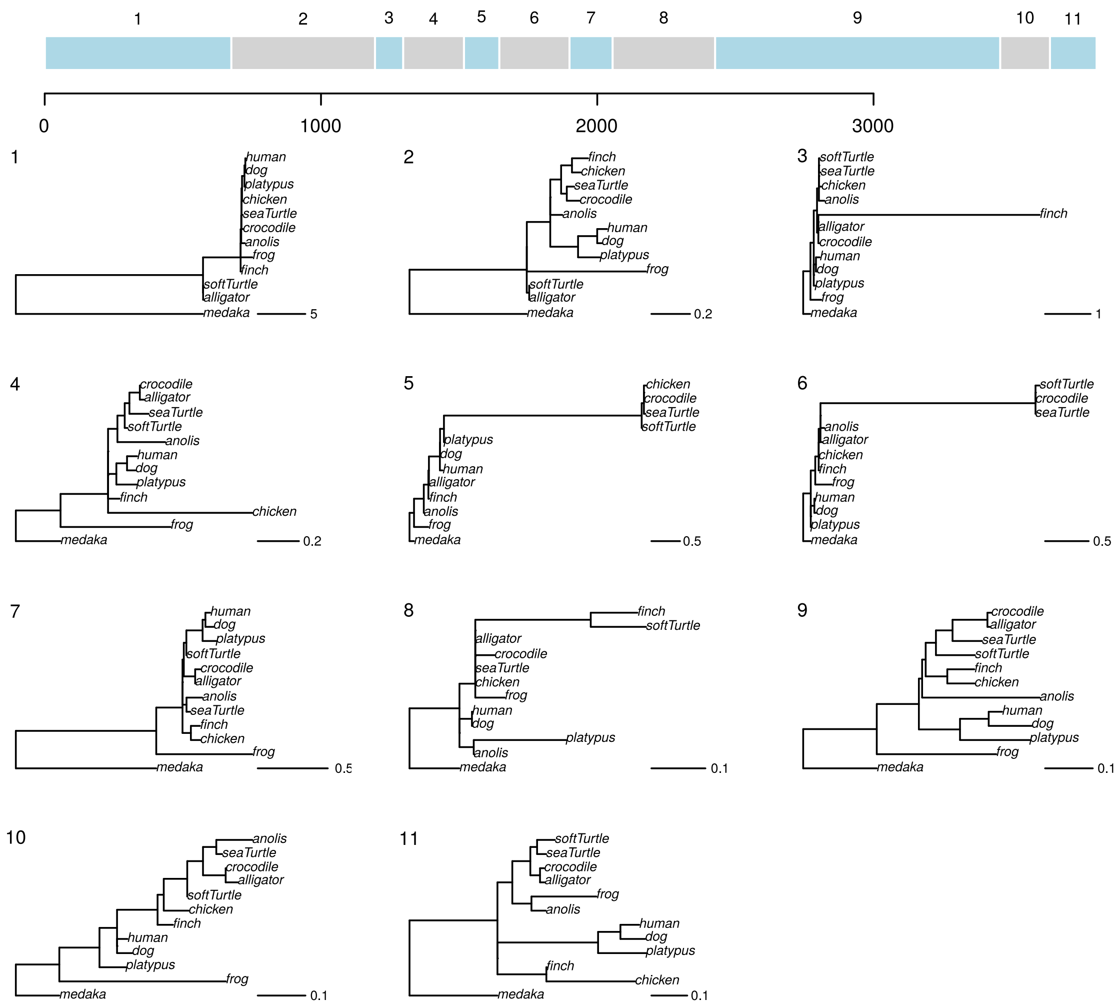


**Figure S40:** replicate three GARD analysis of an FSA realignment of 2798 under GTR+G. Tree topologies correspond to numbered segments, colors are arbitrary. Branch lengths and scale correspond to expected nucleotide substitutions per site.

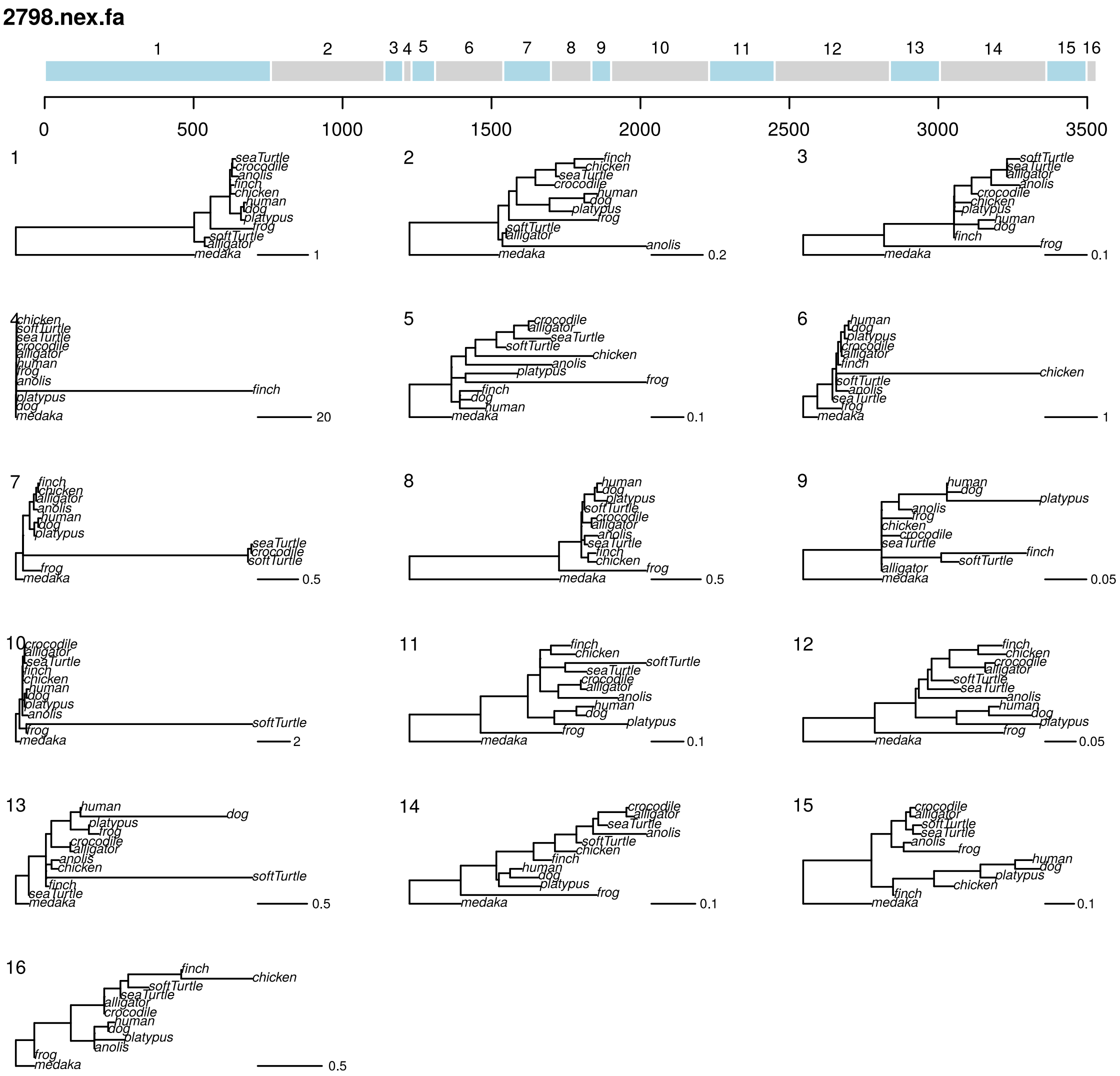


**Figure S41:** replicate one GARD analysis of a PRANK realignment of 2798 under GTR+G. Tree topologies correspond to numbered segments, colors are arbitrary. Branch lengths and scale correspond to expected nucleotide substitutions per site.

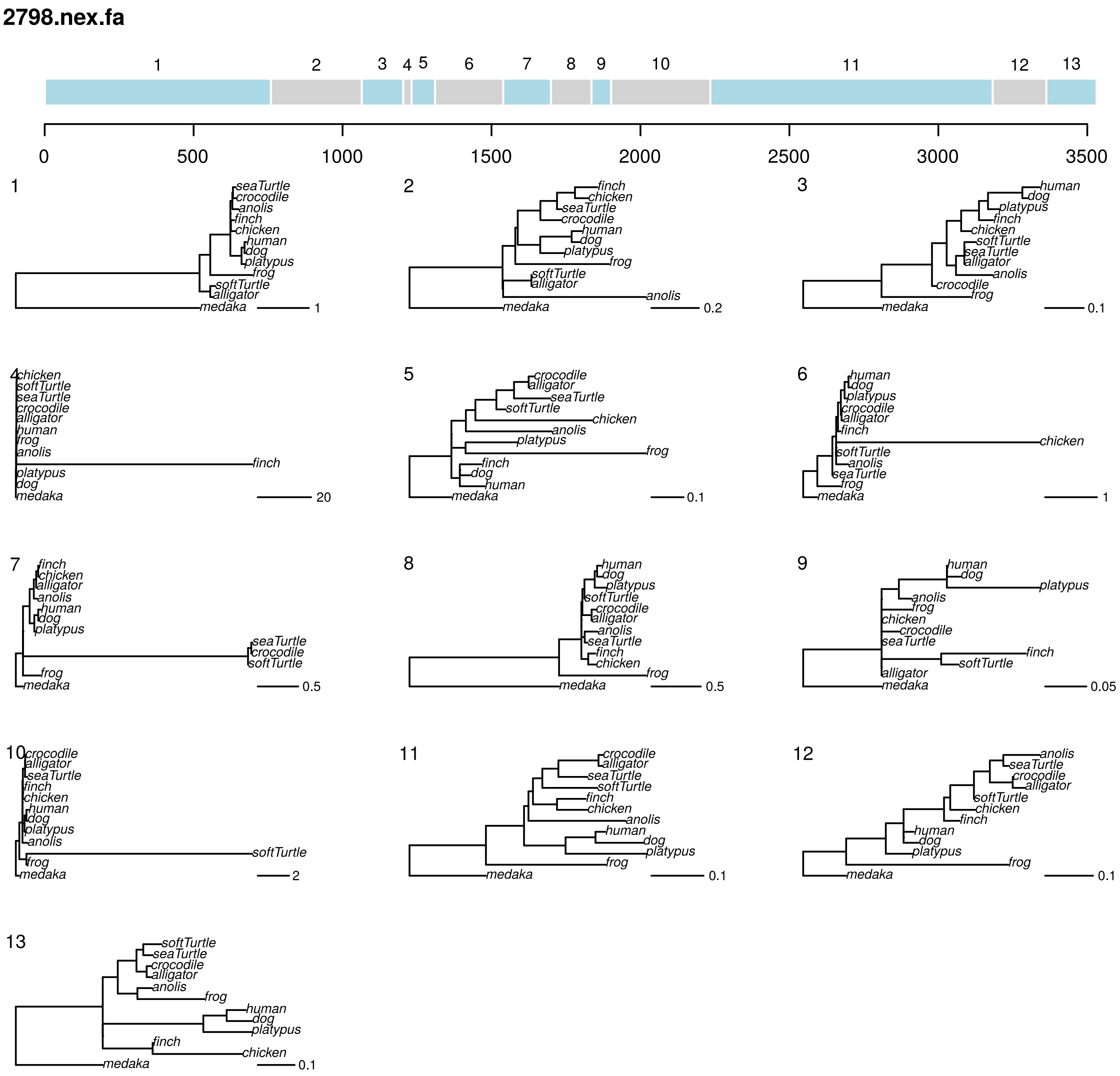


**Figure S42:** replicate two GARD analysis of a PRANK realignment of 2798 under GTR+G. Tree topologies correspond to numbered segments, colors are arbitrary. Branch lengths and scale correspond to expected nucleotide substitutions per site.

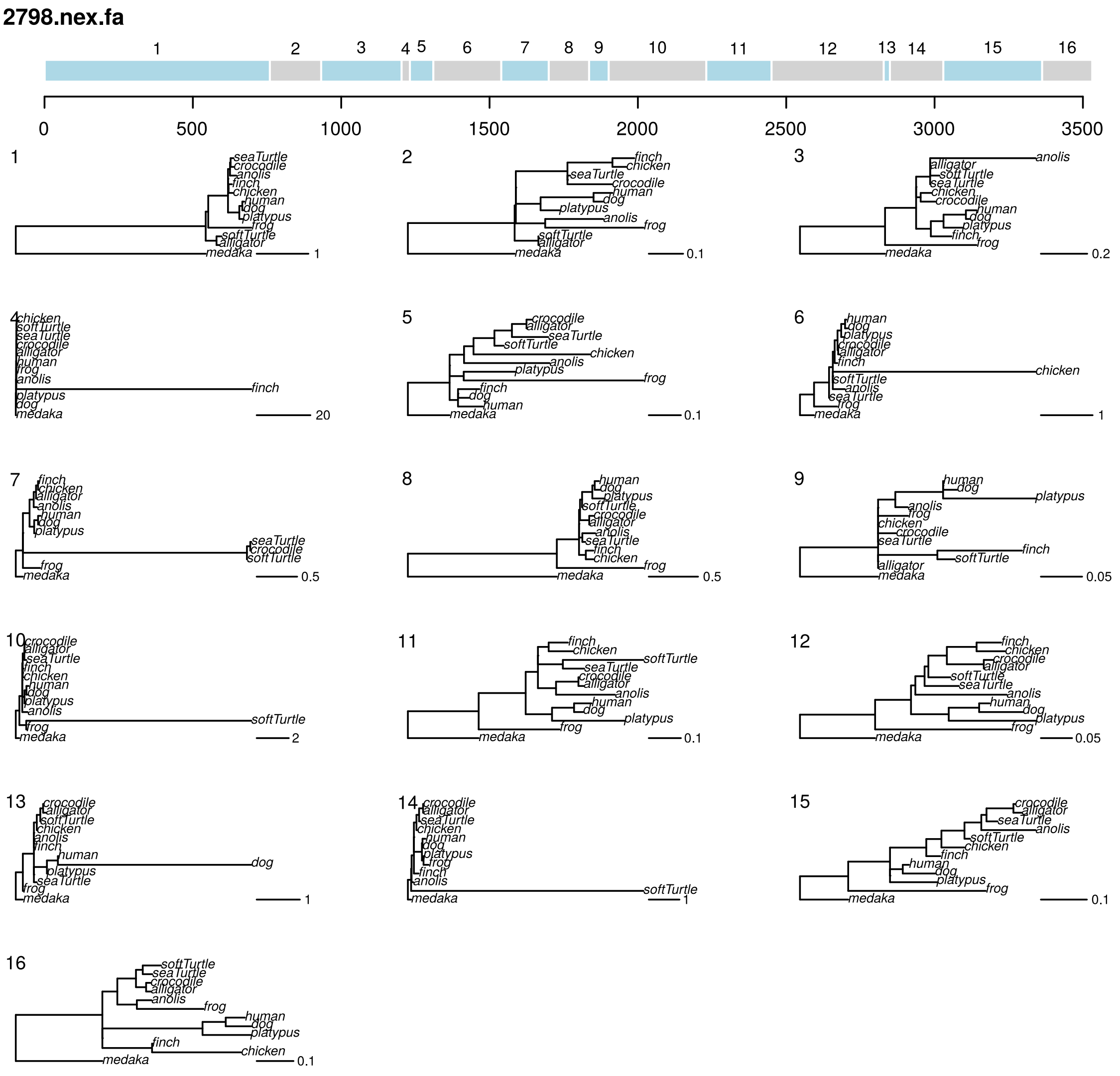


**Figure S43:** replicate three GARD analysis of a PRANK realignment of 2798 under GTR+G. Tree topologies correspond to numbered segments, colors are arbitrary. Branch lengths and scale correspond to expected nucleotide substitutions per site.

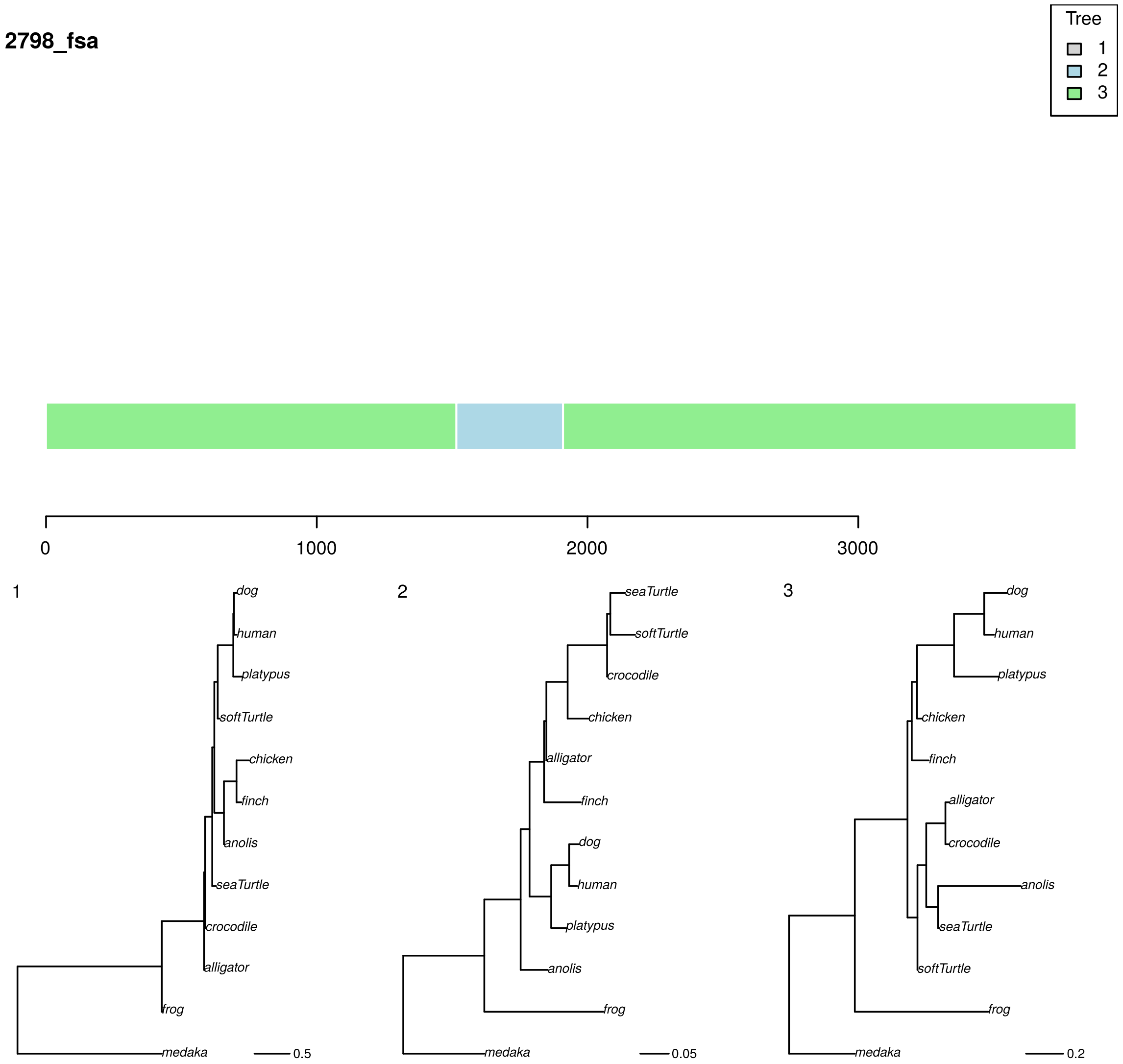


**Figure S44:** phyML_multi analysis of an FSA realignment of 2798 under TN93+G with three trees. Colors correspond to the best-supported tree for each segment. Branch lengths and scale bar correspond to expected nucleotide substitutions per site.

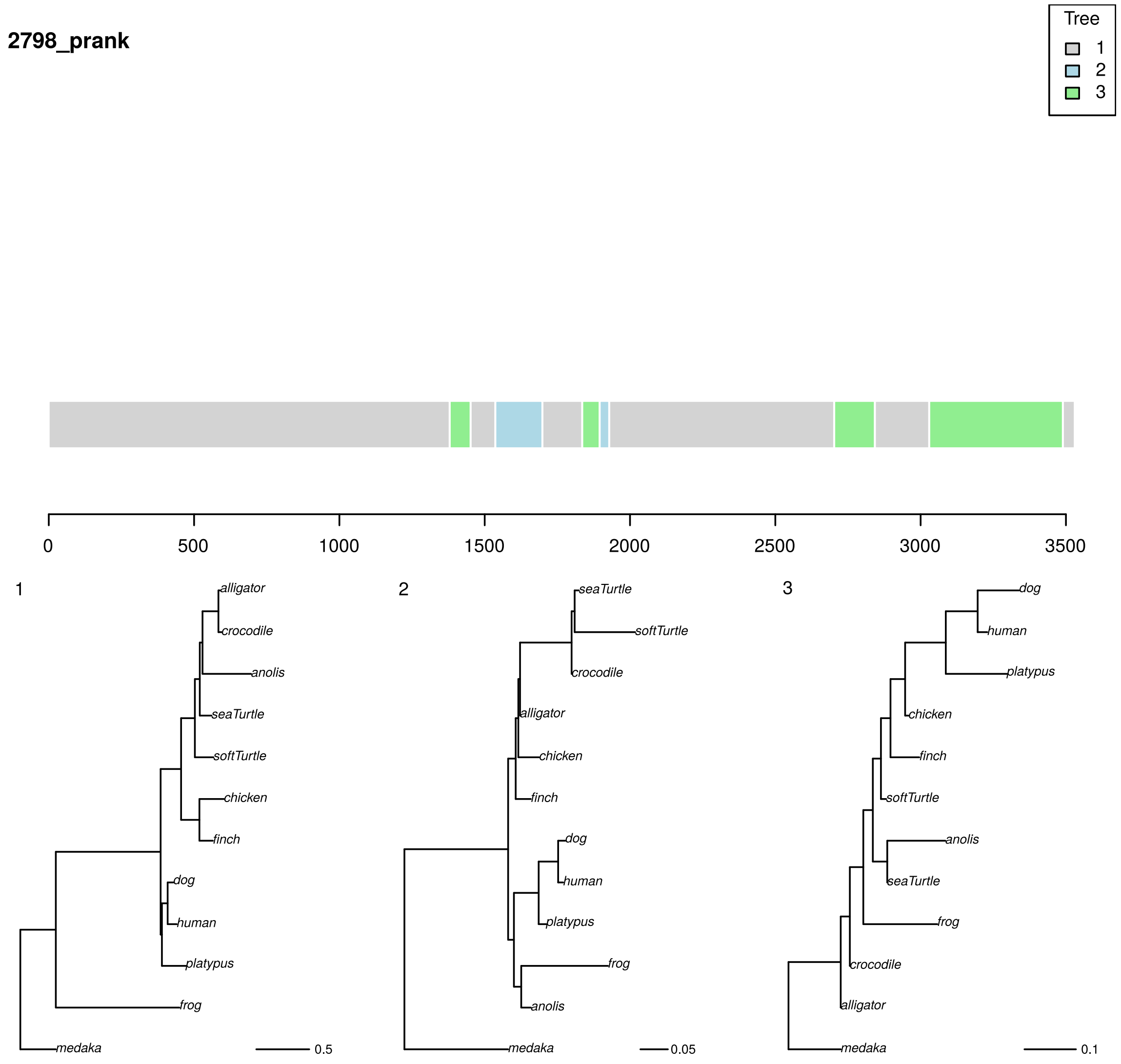


**Figure S45:** phyML_multi analysis of a PRANK realignment of 2798 under TN93+G with three trees. Colors correspond to the best-supported tree for each segment. Branch lengths and scale bar correspond to expected nucleotide substitutions per site.
